# Cytoplasmic Male Sterility and Abortive Seed Traits Generated through Mitochondrial Genome Editing Coupled with Allotopic Expression of *atp1* in Tobacco

**DOI:** 10.1101/2023.06.12.544621

**Authors:** Ralph E. Dewey, Devarshi Selote, H. Carol Griffin, Allison N. Dickey, Derek Jantz, J. Jeff Smith, Anna Matthiadis, Josh Strable, Caitlin Kestell, William A. Smith

## Abstract

Allotopic expression is the term given for the deliberate relocation of gene function from an organellar genome to the nuclear genome. We hypothesized that the allotopic expression of an essential mitochondrial gene using a promoter that expressed efficiently in all cell types except those responsible for male reproduction would yield a cytoplasmic male sterility (CMS) phenotype once the endogenous mitochondrial gene was inactivated via genome editing. To test this, we repurposed the mitochondrially encoded *atp1* gene of tobacco to function in the nucleus under the transcriptional control of a CaMV 35S promoter (construct 35S:nATP1), a promoter that has been shown to be minimally expressed in early stages of anther development. The endogenous *atp1* gene was eliminated (Δ*atp1*) from 35S:nATP1 tobacco plants using custom-designed meganucleases directed to the mitochondria. Vegetative growth of most 35S:nATP1/Δ*atp1* plants appeared normal, but upon flowering produced malformed anthers that failed to shed pollen. When 35S:nATP1/Δ*atp1* plants were cross-pollinated, ovary/capsule development appeared normal, but the vast majority of the resultant seeds were small, largely hollow and failed to germinate, a phenotype akin to the seedless trait known as stenospermocarpy. Characterization of the mitochondrial genomes from three independent Δ*atp1* events suggested that spontaneous recombination over regions of microhomology and substoichiometric shifting were the mechanisms responsible for *atp1* elimination and genome rearrangement in response to exposure to the *atp1*-targeting meganucleases. Should the results reported here in tobacco prove to be translatable to other crop species, then multiple applications of allotopic expression of an essential mitochondrial gene followed by its elimination through genome editing can be envisaged. Depending on the promoter(s) used to drive the allotopic gene, this technology may have potential application in the areas of: (1) CMS trait development for use in hybrid seed production; (2) seedless fruit production; and (3) transgene containment.

## INTRODUCTION

The use of hybrid seeds is critical for maximizing productivity in crop species that show a high degree of heterosis, as they exhibit enhanced growth, fertility and yield compared to their corresponding parental lines (Tester and Langridge, 2010). Traditionally, the most cost-effective hybrid seed systems have exploited a naturally occurring trait known as cytoplasmic male sterility (CMS) to make directional crosses between two inbred lines that display a high degree of hybrid vigor when combined. CMS is the maternally inherited inability to produce viable pollen, and for decades has been successfully used by the seed industry as a facile means for producing commercial-scale quantities of high purity hybrid seed. The maternal inheritance of the CMS trait is attributed to the observation that CMS genes are located within the genome of the mitochondria, an organelle that is strictly inherited through the maternal parent in most angiosperms (Hagemann, 2004).

A classic CMS-based scheme for hybrid seed production involves three lines: (1) the CMS inbred; (2) a maintainer line that is isogenic with the CMS line but contains a normal cytoplasm, enabling it to serve as the pollen parent to maintain and propagate the CMS line; and (3) a restorer line in a different inbred background that possesses dominant ‘restorer of fertility’ (*Rf*) genes that can countermand the CMS trait and enable the F_1_ hybrid plants to be fertile and produce seed when grown by farmers (Chen and Liu, 2014). Examples of crops that have robust CMS-based systems that are used commercially to produce hybrid seed include sorghum, rice, sunflower, and sugar beets. The reasons that CMS-based seed production systems are not applied even more widely are largely due to the absence of efficacious CMS genes, the lack of robust restorer genes, or pleiotropic associations with undesirable traits such as disease susceptibility.

The first CMS gene to be characterized was the *urf13-T* gene of CMS-T maize (Dewey *et al*. 1986; 1987). Molecular investigation of *urf13-T* revealed it to be the product of illegitimate recombination events that combined segments of other mitochondrial genes and genome fragments to yield an open reading frame (ORF) encoding a protein that disrupted male reproductive development, a pattern observed repeatedly as naturally occurring CMS genes from other crop species were elucidated (Chen *et al*. 2017; Gualberto and Newton 2017). The inability to genetically transform plant mitochondria has hindered progress in developing sources of CMS genes beyond those that exist in nature. Because the chloroplast is also maternally inherited, and unlike the mitochondria is amenable to transformation in select plant species, its genome could also be the target for developing novel sources of CMS. In the process of studying the expression of a gene encoding β-ketothiolase in transgenic tobacco chloroplasts, Ruiz and Daniell (2005) made the unexpected discovery that the plants were male sterile. Although this was the first example of an engineered CMS trait, the inability to transform chloroplasts in many crop species, together with the lack of an *Rf* gene that could reverse the sterility in a field environment, greatly limited potential applications of this technology (Chase 2006). Another biotechnology-based approach designed to uncover novel sources of CMS traits involved the RNAi-mediated suppression of the *Msh1* gene whose function has been implicated in the suppression of illegitimate recombination within the mitochondrial genome. Disruption of *Msh1* in both tomato and tobacco resulted in the rearrangement of the mitochondrial genome and/or amplification of preexisting substoichiometric mitochondrial DNA species in a manner that revealed new sources of CMS (Sandhu *et al*. 2007). Similar to the CMS trait generated via chloroplast transformation, however, the utility of CMS genes developed using the *Msh1* inhibition system is limited by the absence of identifiable *Rf* genes to enable seed production in the field.

The advent of genome editing technologies has introduced powerful new tools that can be adapted to modify the mitochondrial genome. Despite having become the system of choice for most genome editing applications, CRISPR/Cas-based technologies are poorly suited for mitochondrial DNA editing, as there is no known reliable method for introducing the guide RNA component across the organelle’s double membrane. By attaching mitochondrial transit peptides to TAL effector nucleases (termed mitoTALENs), it was demonstrated that targeted mitochondrial gene knockout could be achieved within higher plant mitochondria (Kazama *et al*. 2019). In addition to validating the function of two putative CMS genes in rice and rapeseed, this seminal study revealed that double-strand breaks imposed on the mitochondrial genome yielded deletion mutations in the vicinity of the cut sites, and that repair of the broken ends was mediated via homologous recombination across short regions of sequence homology. Since then, mitoTALENs have been used to knock out and validate the function of other proposed CMS genes (Omukai *et al*. 2021; Kuwabara *et al*. 2022; Takatsuka *et al*. 2022), a redundant copy of the essential gene *atp6* in Arabidopsis (Arimura *et al*. 2020), and *nad7*, which encodes a subunit of the NADH dehydrogenase complex (Ayabe *et al*. 2023). Targeted point mutations have also been introduced into the mitochondrial genome by coupling mitoTALENs to a cytidine deaminase base editor (Nakazato *et al*. 2022) or by combining a gene drive-like selection strategy to mitoTALEN-mediated mutagenesis (Forner *et al*. 2022).

Despite the wealth of information that has been accrued with respect to CMS gene function and genome dynamics from the application of mitochondrial-directed genome editing, to our knowledge there have been no reports describing how this technology can be applied to generate traits of agronomic value. We hypothesized that the allotopic expression of an essential mitochondrial gene under the transcriptional control of a promoter that is highly expressed in all tissues except developing anthers, followed by the elimination of the endogenous mitochondrial gene through genome editing, would create a CMS phenotype.

Here, we demonstrate the feasibility of this approach by generating a novel CMS trait in tobacco using *atp1* as the mitochondrial gene target and the 35S *Cauliflower Mosaic Virus* (CaMV) promoter to drive expression of an ATP1 cDNA repurposed to function as a nuclear gene. In addition to the CMS trait, the resultant plants displayed an abortive seed development phenotype upon cross-pollination. Finally, combining the CMS and abortive seed traits within the same plant as described herein may represent the ultimate protection against transgene escape in crop or horticultural species that can be propagated through cuttings or micropropagation.

## RESULTS

### Allotopic expression of an essential mitochondrial gene

We selected the *atp1* gene encoding the α-subunit of the mitochondrial ATPase complex for allotopic expression in tobacco for our proof-of-concept experiments. Our rationale for *atp1* was: (1) unlike most proteins encoded by essential genes within the mitochondria, the Atp1 protein is soluble, and thus less likely to encounter folding issues that may arise as the protein is being imported from the cytosol; and (2) there is precedence for *atp1* functioning properly as a nuclear gene, as it is located in the nuclear genome in animals. Another potential advantage of *atp1* is that the *ATP2* gene that encodes the related β-subunit of the mitochondrial ATPase is a nuclear gene, and thus a favored source for acquiring important elements needed to convert *atp1* into a functional nuclear gene. Specifically, the transit peptide of the *Nicotiana plumbaginafolia* ATP2 protein has been shown to be effective in transporting foreign proteins into the mitochondria (Chaumont *et al*., 1994; Duby *et al*., 2001) and the 3’-untranslated region (UTR) of yeast *ATP2* plays a key role in enhancing ATP2 transport into the mitochondria through mediating the association of *ATP2* transcripts with the organelle (Fox, 2012).

Despite the extensive efforts that have been devoted to the study of plant promoters, we found no promoter that was characterized as being expressed highly in all tissues except those associated with male reproduction. In the absence of such information, we proposed the best candidate to be an enhanced CaMV 35S promoter, given that expression of this promoter was shown to be lacking in the tapetum and sporogenous cells of developing anthers in dicots (Plegt and Bino, 1989). Furthermore, in maize transformed with a glyphosate tolerance gene driven by an enhanced 35S promoter, male reproductive tissues were the only tissue type reported to be unprotected from the herbicide in mature plants sprayed with glyphosate (Feng, *et al.,* 2014).

RNA editing is the phenomenon whereby select nucleotides within mitochondrial transcripts are post-transcriptionally modified (predominantly C to U base transitions), giving rise to mature transcripts whose encoded proteins differ from that predicted by the corresponding DNA sequence (Small, *et al.,* 2020). The tobacco *atp1* gene has six locations where RNA editing has modified codons in the mature transcript (GenBank accession BA000042; Figure S1A). Because of this, when repurposing *atp1* to function within the nucleus, we based the redesign of the gene on the cDNA sequence as opposed to the mitochondrial DNA sequence. Codon usage is another consideration in allotopic expression of mitochondrial genes, as the translational machinery within the organelle is procaryotic-like (Artika, 2020). To avoid the potential of unfavorable codons limiting expression of the repurposed *atp1* gene, a variant was custom synthesized using codons favored in the tobacco nucleus.

The repurposed *atp1* construct, designated 35S:nATP1 (Figure 1A), includes an enhanced 35S promoter upstream of sequences encoding the transit peptide of the tobacco ATP2 protein (including the first 12 aa of the mature protein as these sequences are required for optimal transport; Duby *et al.,* 2001) followed by the codon-optimized *atp1* cDNA. The presumed 3’-UTR and termination signal of the tobacco *ATP2* gene were also used. 35S:nATP1 (nucleotide sequence, Figure S1B) was cloned into expression vector pCAMBIA1300 which contains the *hptII* selectable marker gene.

**Figure 1.**
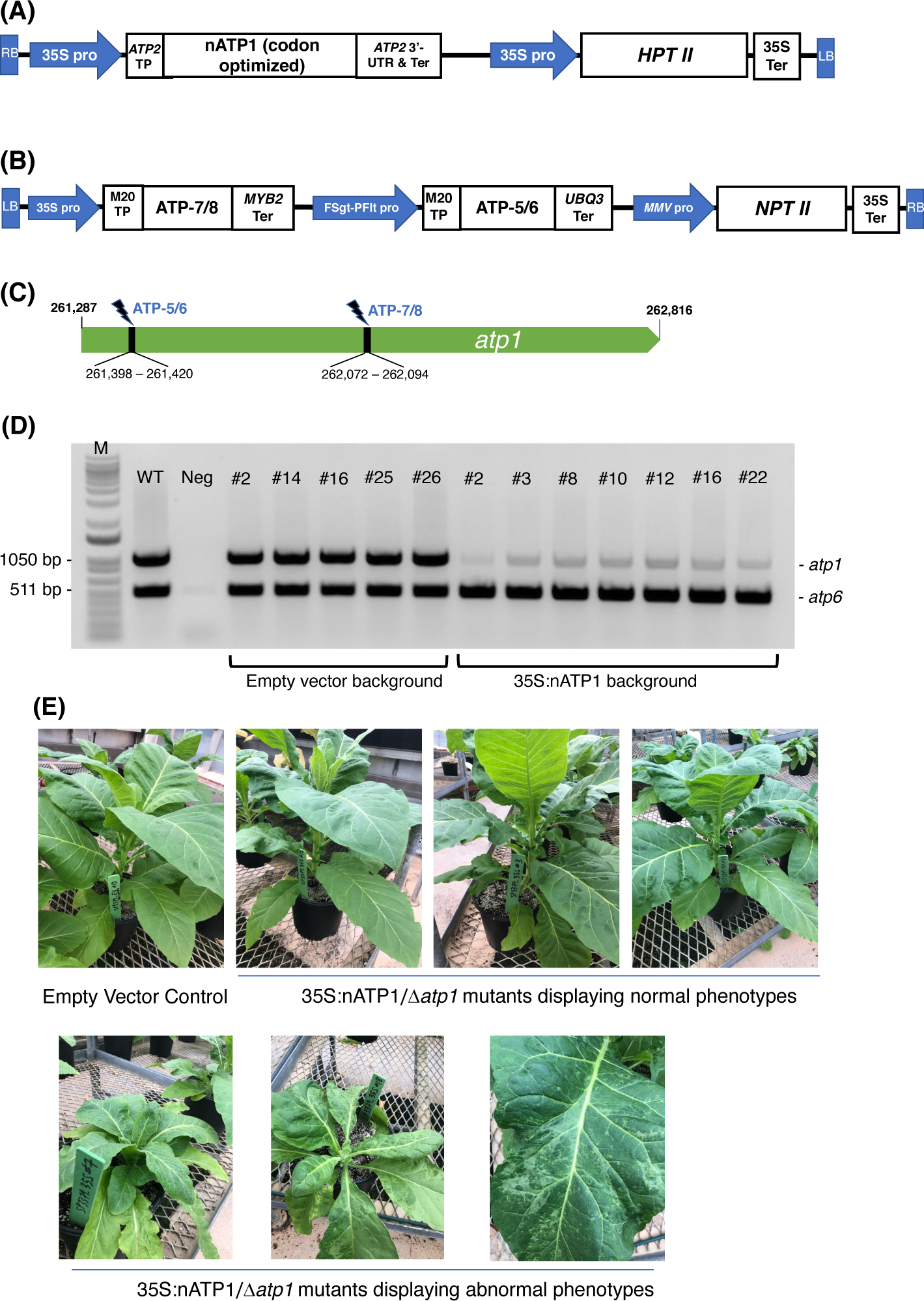
Mitochondrial genome editing of *atp1* in 35S:nATP1 transformed plants. **(A)** 35S:nATP1 construct. LB, left border; RB, right border; 35S pro, CaMV 35S promoter; TP, transit peptide; UTR, untranslated region; Ter, terminator; *HPTII*, *hygromycin phosphotransferase II*. **(B)** *atp1-*targeting vector SP3379 containing mitoARCUS cassettes ATP-7/8 and ATP-5/6. FSgt-PFlt pro, *Figwort Mosaic Virus – Peanut Chlorotic Streak Caulimovirus* chimeric promoter; MMV pro, *Mirabilis Mosaic Virus* promoter; *NPTII*, *neomycin phosphotransferase II*. **(C)** Tobacco *atp1* gene. MitoARCUS cleavage sites are indicated with thunderbolts. Nucleotide positions are in accordance to the tobacco mitochondrial reference genome BA000042. **(D)** PCR analysis of individual T0 plants transformed with SP3379. Each reaction contained primers designed to simultaneously amplify *atp1*, and *atp6* as a control. M, molecular size markers; WT, wild type K326 tobacco; Neg, negative control lacking template DNA. **(E)** T0 35S:nATP1 plants lacking a functional mitochondrial *atp1* gene (Δ*atp1*) six weeks after transplanting to soil. Phenotypically normal plants are shown in the upper panel; individuals with aberrant leaf or growth phenotypes are shown in the lower panel. K326 EV#13, empty vector control plant.

Transgenic plants were generated by transforming haploid plants of tobacco variety K326 with the 35S:nATP1 construct. Eighteen hygromycin resistant T_0_ individuals were screened for transgene expression using semi-quantitative RT-PCR (Figure S2). Individuals judged to be expressing nATP1 to the greatest extent were chromosome doubled to produce a fixed, high expressing nATP1 line (35S:nATP1#25) used for *atp1* editing.

### *atp1* genome editing

Custom-designed meganuclease enzymes developed via a genome editing platform termed ARCUS (Precision BioSciences) were used to target and cleave *atp1*. ARCUS enzymes are created by protein engineering of the homing endonuclease I-CreI and have been successfully designed to target and cleave the mitochondrial genome of mice (termed mitoARCUS; Zekontye *et al*., 2021). I-CreI exists as a dimer, but mitoARCUS enzymes contain an engineered linker that fuses the two engineered monomers into a single polypeptide. One unique property that distinguishes ARCUS enzymes from other genome editing platforms is the production of 4 bp 3’-overhangs upon cleavage. This presented a unique opportunity to test whether precise deletions in the mitochondrial genome could be created via the cleavage and ligation of compatible sticky ends by two mitoARCUS enzymes targeting sequences in close proximity, as opposed to the somewhat random deletions observed by researchers using mitoTALENs. A single plant transformation vector, designated SP3379, was developed with two distinct ARCUS cassettes (ATP-5/6 and ATP-7/8) designed to target the tobacco *atp1* gene (Figure 1B and 1C). SP3379 possesses an *nptII* gene to enable kanamycin selection (Figure S3). Both ARCUS enzymes were predicted to leave the same 4 bp 3’-overhang following cleavage of their respective target sites (Figure S1A), the re-ligation of which would result in a 674 bp deletion. To facilitate entry of the ARCUS enzymes into the mitochondria, sequences encoding a transit peptide designated M20 were introduced immediately upstream of those encoding the ARCUS proteins, as this transit peptide was shown to be effective in transporting ARCUS-GFP fusions into the mitochondria of tobacco protoplasts (Figure S4).

Doubled haploid K326 line 35S:nATP1#25 was transformed with vector SP3379. As a control, leaf discs from an empty vector (EV) K326 line containing only the pCAMBIA1300 vector were also transformed. Fifteen T_0_ plants were recovered from the 35S:nATP1/SP3379 transformation experiment and nine plants were recovered from K326 EV transformed with SP3379. To test whether *atp1* had been altered in any of the plants transformed with the mitoARCUS constructs, PCR primers were designed corresponding to *atp1* sequences flanking both the ATP-5/6 and ATP-7/8 recognition sites. As a control, primers specific for the mitochondrial gene *atp6* were also included in each PCR reaction. Remarkably, in all 15 T_0_ plants in the 35S:nATP1 background the intensity of the *atp1* amplification product was greatly reduced in comparison to *atp6*; in contrast, all nine plants recovered in the K326 EV line showed *atp1* and *atp6* amplification products of similar intensity. In no instance was an *atp1* amplification product observed that possessed a 674 bp deletion indicative of the perfect removal and re-ligation of the sequences between the ATP-5/6 and ATP-7/8 target sites.

There are two plausible explanations for the observation of *atp1* amplification products being greatly diminished, but not fully eliminated in 35S:nATP1/SP3379 plants: (1) the cleavage of *atp1* yielded heterogeneous populations of mutated and WT *atp1* genes; or (2) the minor amplification products in these plants are false positives, corresponding to <u>nu</u>clear-encoded <u>mit</u>ochondrial DNA sequence<u>s</u> (NUMTs). Over the course of evolution, large portions of the mitochondrial genome have become incorporated into the nuclear genome (Lough *et al.,* 2015; Yoshida *et al.,* 2017). Although these presumably nonfunctional sequences have accumulated polymorphisms over time, it can be difficult to differentiate a true mitochondrial gene from an NUMT using PCR-based assays (Arimura *et al*. 2020).

As an initial attempt to discriminate between true *atp1* sequences and *atp1*-like NUMTs, we sought to design more specific primers based on NUMTs found in the tobacco genome. Utilizing BLASTN searches on draft genome sequences of *N. tabacum* with *atp1* as the query sequence, over 30 scaffolds or contigs were found that shared greater than 93% sequence identity to the *atp1* sequence. Three forward and three reverse primers were designed based on the NUMT information and tested for their ability to amplify *atp1* in 35S:nATP1/SP3379 events. Although considerable variability was observed in *atp1* band intensity among the different primer combinations, in no case was the complete absence of a band observed (Figure S5). For subsequent experiments, we typically used the primer pair that regularly yielded the faintest PCR product (atp1_F3 and atp1_R2; Table S1). PCR analysis of seven 35S:nATP1/SP3379 events and five EV control plants using this primer pair is shown in Figure 1D.

As a direct test of the nature of the faint amplification product observed in 35S:nATP1/SP3379 plants, the PCR products generated by one of the *atp1* primer pairs (atp1_F1 and atp1_R1) from two 35S:nATP1/SP3379 events (#8 and #16) and a WT K326 control were cloned into a plasmid vector. Sequencing runs were conducted on 22 – 24 independent plasmids from each of the three cloning experiments (Data S1 – S3). None of the amplification products from plants 35S:nATP1/SP3379#8 or 35S:nATP1/SP3379#16 (hereafter referred to as 35S:nATP1/Δ*atp1*#8 and 35S:nATP1/Δ*atp1*#16, respectively) were 100% identical to the WT *atp1* sequence. Instead, each contained polymorphisms that could be found in one or more *atp1*-like NUMT (Data S4). Interestingly, there were no NUMTs on a tobacco genomic contig that perfectly coincided across the entire read of a given PCR amplification product, suggesting that additional *atp1*-like NUMTs exist in the tobacco genome that are not present in current drafts and/or the existing tobacco contigs were not assembled accurately in these regions. In contrast, 21 of 24 cloned PCR products amplified from the WT control were identical to *atp1*. These results suggest that an intact *atp1* gene is no longer present in the mitochondrial genomes of plants expressing mitoARCUS enzymes. To gain insights into the nature of the nine plants in the K326 EV background that were recovered, we assayed these for the presence of the *nptII* selectable marker gene. Seven of the nine plants were negative for *nptII*, suggesting they were escapes. It is likely that the mitoARCUS cassettes were minimally expressed in the two *nptII-*positive plants.

### Phenotypic evaluation of 35S:nATP1/Δ*atp1* plants

T_0_ plants were transferred to soil and grown indoors until 10 – 15 cm in height, then transferred to large pots and grown to maturity in a greenhouse. Nine of the fifteen 35S:nATP1/Δ*atp1* plants appeared phenotypically normal during vegetative growth. The other six, however, displayed a reduced growth and/or a misshapen leaf phenotype with mosaic patterning (Figure 1E). By ten weeks after transplant, the phenotypically normal 35S:nATP1/Δ*atp1* individuals had grown to a typical height and had flowered, while the abnormal plants continued to show a slow growth phenotype (Figure S6). The first distinguishing characteristic between otherwise normal looking 35S:nATP1/Δ*atp1* plants and WT (or EV control) plants became apparent upon flower formation. At full maturity (Stage 12 as defined by Koltunow *et al*. 1990) the petals of 35S:nATP1/Δ*atp1* individuals were less expanded at the top (Figures 2A and S7A). Examination of the anthers revealed even more dramatic differences. The anthers of 35S:nATP1/Δ*atp1* plants appeared malformed and failed to dehisce (Figures 2B and S7B), while the pistils and stigmas appeared normal. Pollen staining within intact Stage 10 anthers, followed by their release by pressing the coverslip firmly on the slide, revealed plentiful pollen in WT tobacco (both free and still associated with the anther). In contrast, no viable pollen grains were detected in the 35S:nATP1/Δ*atp1* anthers (Figure 2C).

**Figure 2.**
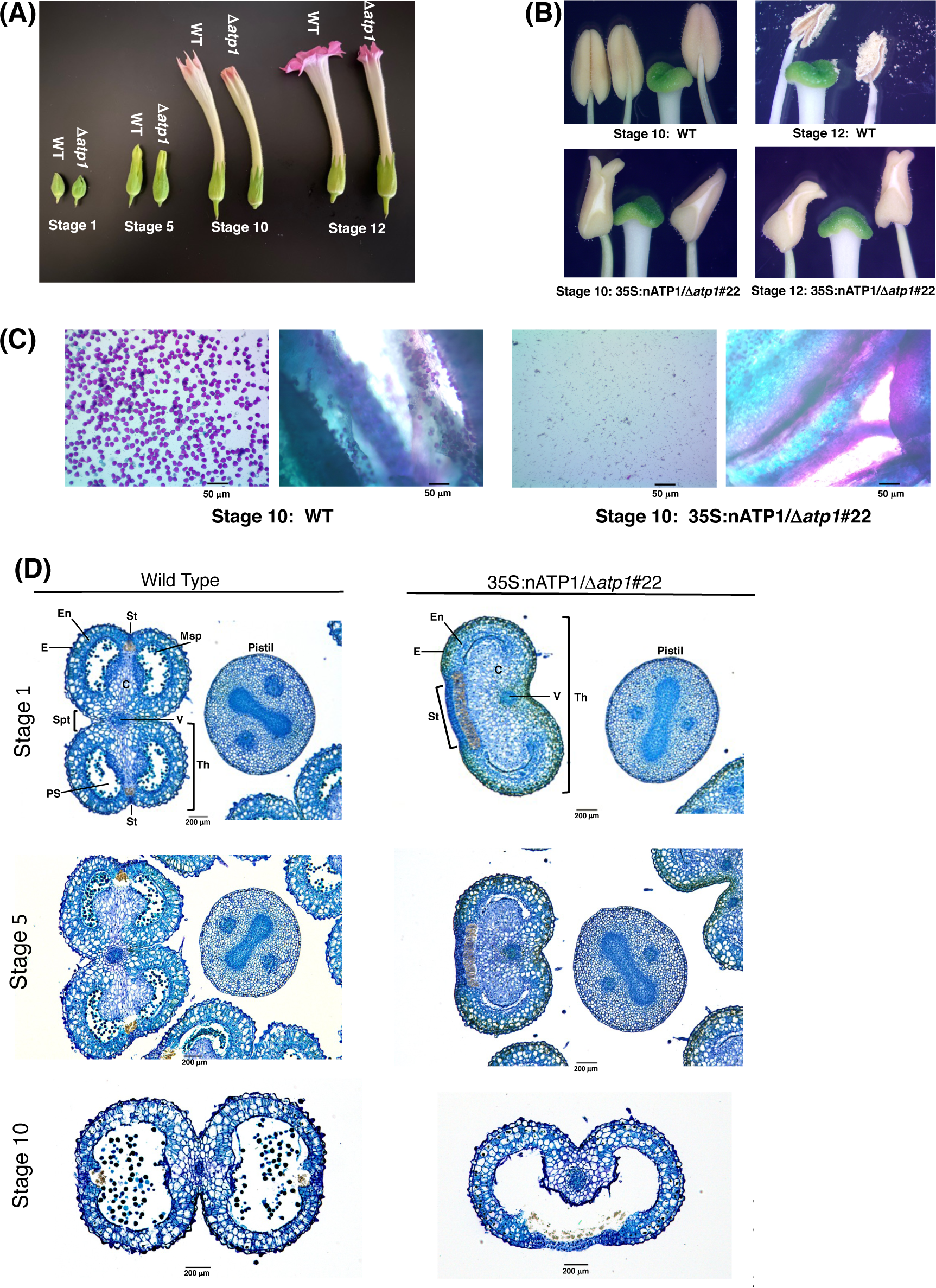
35S:nATP1/Δ*atp1* plants have abnormal flowers, misshapen anthers, and fail to produce pollen. **(A)** Flower development in K326 WT and 35S:nATP1/Δ*atp1*#22 (Δ*atp1*) individuals at Stages 1, 5, 10 and 12 (as defined in Koltunow *et al*. 1990). **(B)** Anthers and stigmas at Stages 10 and 12 in K326 WT and 35S:nATP1/Δ*atp1*#22 mutant plants. **(C)** Staining of Stage 10 anthers from K326 WT and 35S:nATP1/Δ*atp1*#22 individuals. For each genotype, the panel on the left shows the contents expelled upon firmly pressing the stained anthers on a microscope slide with a cover slip. Viable pollen grains are stained purple. Panels on the right show a cross section of the anthers after pollen expulsion. **(D)** Histological analysis of Stage 1, Stage 5 and Stage 10 anthers. C, connective tissue; E, epidermis; En, endothecium; Msp, microspores; PS, pollen sac; St, stomium; Spt, septum; Th, theca; V, vascular bundle. Scale bars are indicated.

Toluidine Blue-O staining of paraffin-embedded histological sections provided a more detailed analysis of the differences between normal anthers and those observed in 35S:nATP1/Δ*atp1* plants. WT tobacco anthers contain two distinct halves, or theca, separated by the vascular bundle. 35S:nATP1/Δ*atp1* anthers contained a single kidney-shaped theca (Figure 2D). At Stage 1, microspores had already formed in pollen sacs of the two locules within each theca of WT anthers; in contrast, pollen sacs were collapsed with no evidence of microspore formation in 35S:nATP1/Δ*atp1* anthers. At Stage 5, pollen sacs had become apparent in the mutant anthers, but were lacking microspores. At Stage 10, the two locules within each theca had fused and were filled with pollen in the WT anthers. The two locules had similarly fused within the single theca of the mutant anthers by Stage 10, with still no evidence of pollen grains. A morphological distinction that was apparent throughout anther development was the appearance of an exaggerated stomium region in 35S:nATP1/Δ*atp1* anthers. The enlarged stomium is apparent both on the histological sections (Figure 2D) and close-up pictures of the mutant anthers (the whitish triangular sections of the 35S:nATP1/Δ*atp1* anthers in Figure 2B). The stomium is the specialized cell layer of the anther that ruptures upon dehiscence, facilitating the release of mature pollen grains (Keijzer 1987). Despite its exaggerated structure, the stomium in Δ*atp1* mutant anthers remained intact throughout anther maturation. Collectively, these results suggest that 35S:nATP1/Δ*atp1* plants are male-sterile, and because the perturbation is attributable to a mutation in the mitochondrial genome, they therefore manifest a CMS phenotype.

To propagate the 35S:nATP1/Δ*atp1* CMS materials, 35S:nATP1 and WT K326 plants were used as pollen parents in crosses to the T_0_ individuals. Minimal evidence of ovary development was observed in unfertilized flowers of 35S:nATP1/Δ*atp1* plants (Figures 3A, S6C, and S8A). In contrast, ovary development and capsule formation in fertilized 35S:nATP1/Δ*atp1* individuals appeared similar to that observed in self-fertilized WT and EV controls (tobacco is typically a self-fertilizing species). Subtle differences were observed, however, between the seed collected from 35S:nATP (or WT) X 35S:nATP1/Δ*atp1* crosses compared to the controls, as seed from the crosses appeared to be both smaller and lighter in color (Figures 3B and S8B). One-hundred seed weight counts revealed that seeds from WT and 35S:nATP plants weighed 7.3 to 14.4 times more than seeds from the crosses (Figure 3C). In total, crosses were made between 35S:nATP1 and/or WT K326 pollen parents and eight of the 35S:nATP1/Δ*atp1* T_0_ individuals (35S:nATP1/Δ*atp1*#1, #3, #8, #11, #12, #21, #22, and #23). When these seeds were placed on soil no germination was observed.

**Figure 3.**
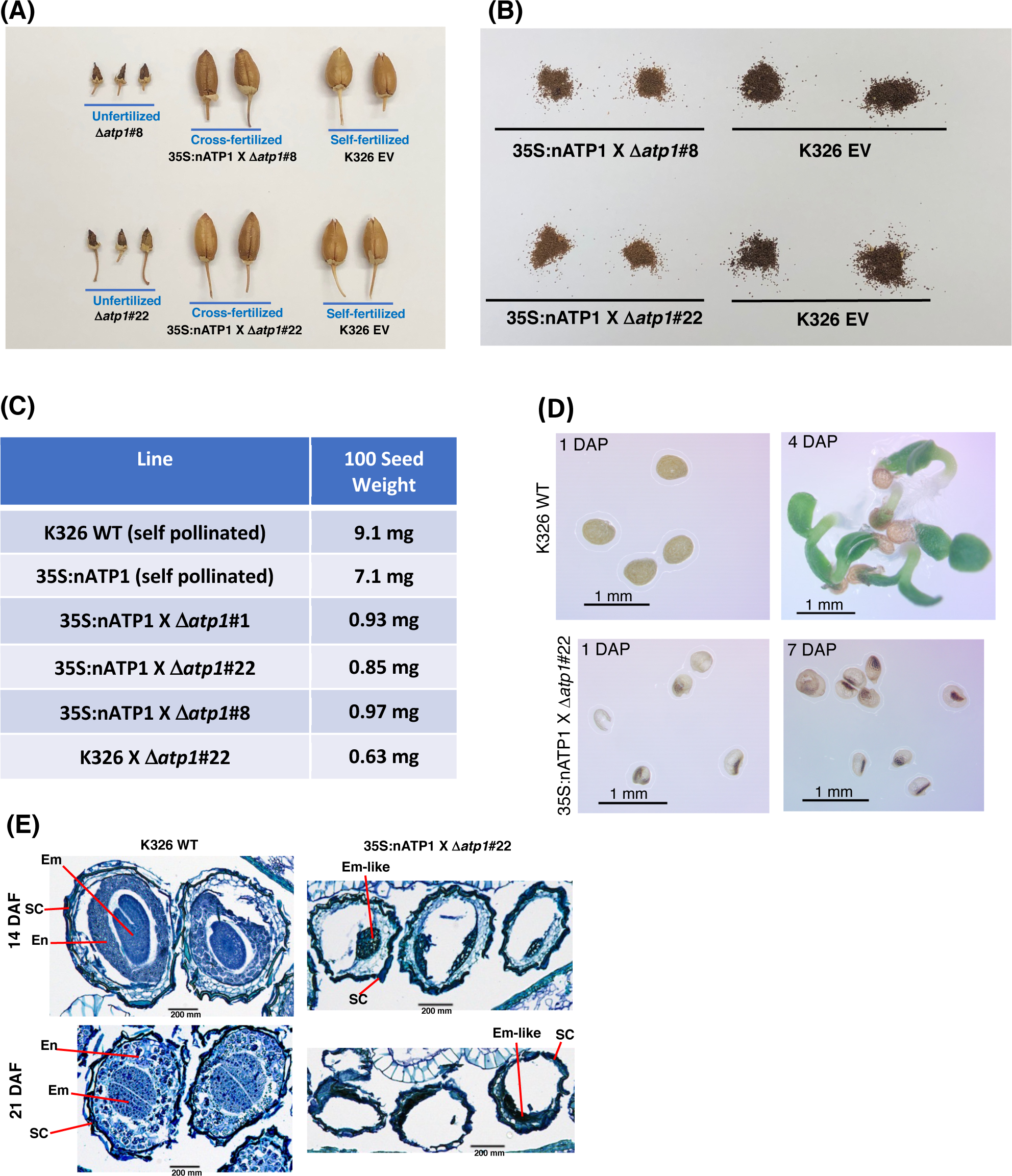
Seed development is aborted in cross-fertilized 35S:nATP1/Δ*atp1* plants. **(A)** Capsules fail to develop in unfertilized 35S:nATP1/Δ*atp1*#8 (Δ*atp1*#8) and 35S:nATP1/Δ*atp1*#22 (Δ*atp1*#22) flowers, but appear normal when crossed to a 35S:nATP1 pollen parent. (B) Seeds produced from 35S:nATP1 X 35S:nATP1/Δ*atp1* crosses are smaller and lighter in color than EV control tobacco seeds. (C) Hundred-seed weights of 35S:nATP1/Δ*atp1*#1 and #22 plants fertilized with 35S:nATP1 pollen. (D) 35S:nATP1 X 35S:nATP1/Δ*atp1* seeds are translucent and fail to germinate on solid MS media. DAP, days-after-plating. (E) Stained histological sections of developing WT and 35S:nATP1 X 35S:nATP1/Δ*atp1* seeds at 14 and 21 days-after-fertilization (DAF). Em, embryo; En, endosperm; SC, seed coat. Scale bars are indicated.

To better observe the differences between WT seeds and those obtained from 35S:nATP X 35S:nATP1/Δ*atp1* crosses, seeds were germinated on solid MS media. At one day-after-plating (DAP), WT seeds were opaque. Seeds from 35S:nATP X 35S:nATP1/Δ*atp1* crosses, however, were smaller and translucent, with most of the seeds containing a small, dense interior region (Figure 3D). By four DAP, small plantlets had emerged from the WT seeds, but even at seven DAP the seeds from the crosses appeared no different than those at one DAP. Although several hundred seeds from 35S:nATP X 35S:nATP1/Δ*atp1* crosses were plated on MS media, no germination was observed. Histological staining was also conducted during seed development.

At 14 days-after-fertilization (DAF), torpedo-shaped embryos were visible, surrounded by layers of developing endosperm cells in self-fertilized WT plants (Figure 3E). By 21 DAF, when tobacco seeds are fully mature, there were very few non-stainable sectors within the WT tobacco seeds. In contrast, at both 14 and 21 DAF, the majority of the volume within the developing 35S:nATP X 35S:nATP1/Δ*atp1* seeds remained unstained, and there was little evidence of endosperm development. At both stages there were clusters of darkly staining cells that we speculate to be embryo-like structures.

### Characterization of mitochondrial genomes in 35S:nATP1/Δ*atp1* plants

Due to the inability to perpetuate Δ*atp1* T_0_ events through seed, tissue culturing of leaf discs followed by plant regeneration was conducted to propagate these materials. Three tissue culture-maintained lines were selected for PacBio-based sequence analysis: 35S:nATP1/Δ*atp1*#8, 35S:nATP1/Δ*atp1*#16, and 35S:nATP1/Δ*atp1*#22. Lines 35S:nATP1Δ/*atp1*#8 and #22 were selected as representatives of T_0_ events that displayed normal growth habits while 35S:nATP1Δ*atp1*#16 was chosen from among the events noted for slow growth and short stature, a phenotype observed in both the original T_0_ plant as well as subsequent plants derived from tissue culture. A picture of tissue culture-propagated plants of similar age representative of the three selected lines is shown in Figure S9.

Due to the large size of the tobacco genome (4.5 Gb), ground leaf preparations were subjected to several rounds of differential centrifugation to enrich the crude fractions for mitochondria prior to DNA extraction for sequencing. PacBio sequencing of the DNAs isolated from 35S:nATP1/Δ*atp1*#8, #16, and #22 yielded read counts of 878,706, 543,041, and 911,783, respectively (Table S2). Of these, 3.4 to 4.9% displayed sequence homology to the tobacco mitochondrial reference genome BA000042, which included an assortment of legitimate mitochondrial DNA reads, NUMTs, and even chloroplast DNA, as fragments of the chloroplast genome are scattered throughout plant mitochondrial genomes (Kubo and Newton 2008). To enhance for true mitochondrial DNAs, reads that were more than 33% clipped were removed prior to visualizing read coverage to the reference genome. A gap was observed in the region of *atp1* in all three 35S:nATP1/Δ*atp1* lines when reads were mapped to BA000042 (Figures 4A-C).

**Figure 4.**
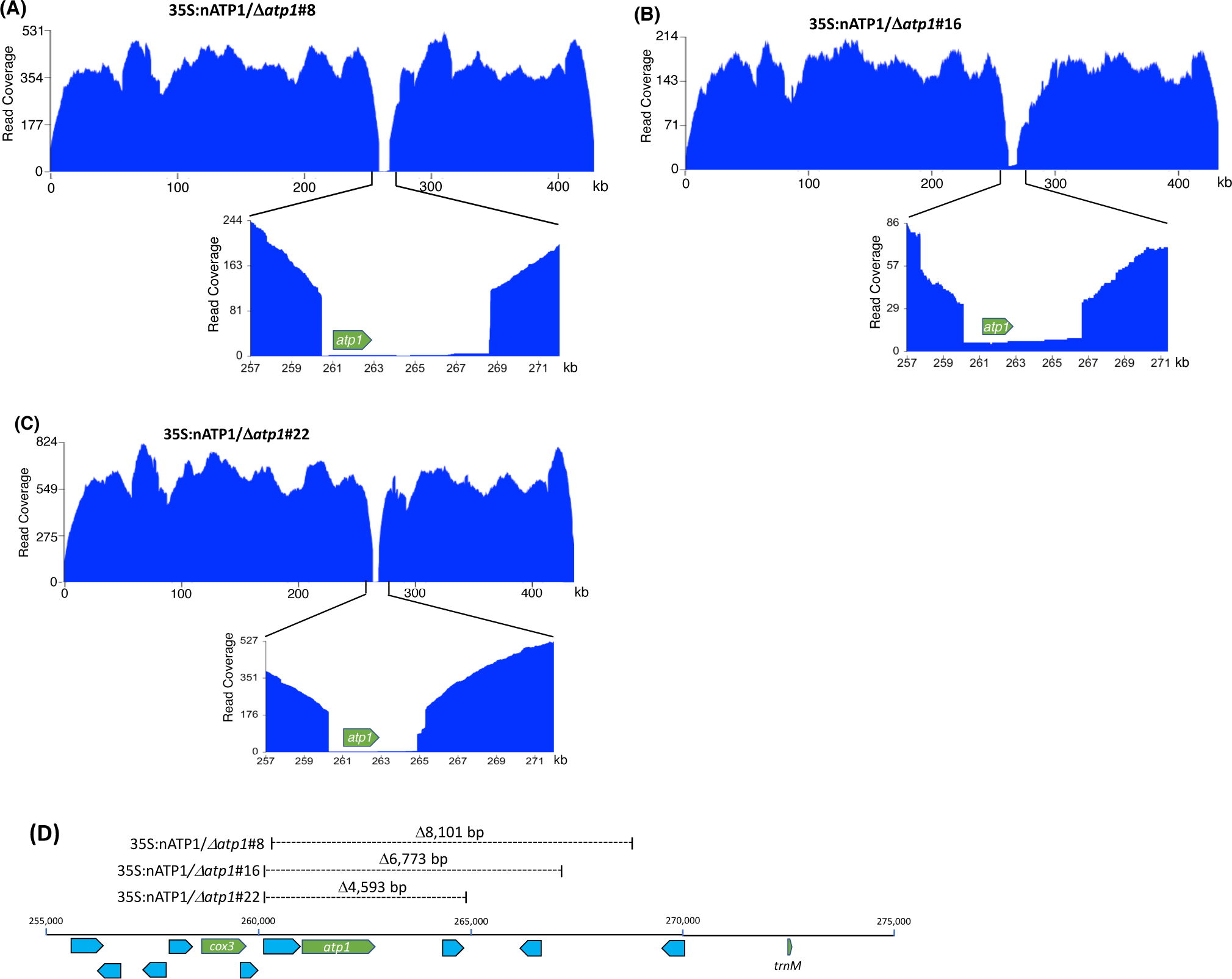
Long read sequence coverage of the mitochondrial genomes of 35S:nATP1/Δ*atp1*#8 (A), #16 (B), and #22 (C). x-axis, reference genome BA000042; y-axis, PacBio sequence coverage after removal of 33% clipped reads. **(D)** Position and extent of sequences missing in Δ*atp1* mutants. Known functional genes *cox3, atp1* and *trnM* are highlighted in green; uncharacterized ORFs greater than 100 codons in length are highlighted blue.

Despite the reads being greatly diminished in the region of *atp1*, a small number did span across the entire reading frame in 35S:nATP1/Δ*atp1*#8 (n=5), #16 (n=7), and #22 (n=4) mitochondria. Alignment of the ORFs corresponding to *atp1* from each of these reads (Data S5) to the WT *atp1* sequence revealed polymorphisms that were shared with NUMTs predicted to be located in the nuclear genome (Data S4) as well as the PCR products of the low representation *atp1-like* sequences observed when events 35S:nATP1/Δ*atp1*#8 and #16 were amplified using primers intended to be specific for *atp1* (Data S2 and S3). In general, the PacBio *atp1*-like NUMTs matched more closely to the PCR products than the NUMTs in GenBank, again suggesting that these regions of the tobacco draft genome are incomplete or inaccurately assembled. The PacBio-based NUMT that shared the highest sequence homology to WT *atp1* differed only by three polymorphisms. Four of the PCR products amplified from event 35S:nATP1/Δ*atp1*#8 (Data S2) and two PCR products from event 35S:nATP1/Δ*atp1*#16 (Data S3) corresponded perfectly with this PacBio NUMT.

The genuine mitochondrial-specific PacBio reads assembled into five predicted contigs in lines 35S:nATP1/Δ*atp1*#8 and #22, and four predicted contigs in line 35S:nATP1/Δ*atp1*#16 (Table S2). Circos plots aligning the predicted contigs to BA000042 showed that collectively, every region of the WT reference genome was represented in the predicted contigs, except the region surrounding *atp1* (Figure S10). Examination of the contigs in these areas revealed predicted 5’ recombination junctions in the region between the *cox3* gene and *atp1*, and 3’ recombination junctions in the region between *atp1* and a *trnM* gene (Figure 4D). Specifically, the mitochondrial genomes of lines 35S:nATP1/Δ*atp1*#8, #16, and #22 were predicted to harbor deletions of 8,101 bp, 6,773 bp, and 4,593 bp, respectively. Although a small number of uncharacterized ORFs are found in this region, no gene of known function other than *atp1* is found in the sequences predicted to be deleted. Because the extent of the deletion observed in line 35S:nATP1/Δ*atp1*#16 was smaller than that found in 35S:nATP1/Δ*atp1*#8, the slow growth phenotype of the former cannot be simply explained by a larger proportion of the mitochondrial genome having been deleted.

Close investigation of the predicted recombination junctions in the Δ*atp1* mutant genomes revealed regions of homology ranging from 6 bp to 188 bp (Figure 5). For line 35S:nATP1/Δ*atp1*#8, a 188 bp sequence between *cox3* and *atp1* appeared to have recombined with a homologous 188 bp sequence (differing by only 3 polymorphisms) found over 57 kb away in reverse orientation (Figure 5B). Downstream of *atp1*, a recombination event between an inverted 9 bp repeat sequence separated by 55 bp was predicted to result in a large 12,719 bp inverted repeat. Events 35S:nATP1/Δ*atp1*#16 and #22 shared the same predicted recombination event in the region 5’ of *atp1* (Figure 5C and 5D). A 148 bp sequence between *cox3* and *atp1* is 100% identical to a 148 bp sequence, in reverse orientation, found in the middle of the large 4,713 bp Repeat 2 (Rep2) that is located three times on the master reference genome (Figure 5A). From the sequence configurations predicted in the contigs of lines 35S:nATP1/Δ*atp1*#16 and #22, as well as alignments of individual PacBio reads, it appeared that recombination could occur between the 148 bp sequence upstream of *atp1* and any of the three Rep2 copies in the reference genome. The recombination event predicted in the region 3’ of *atp1* in line 35S:nATP1/Δ*atp1*#16 appeared to have occurred across a 6 bp sequence of microhomology (ACCACC) between sequences located over 32 kb away (Figure 5c). Finally, the 3’ recombination event in line 35S:nATP1/Δ*atp1*#22 was predicted to have occurred across an 8 bp inverted repeat separated by 421 bp, giving rise to a large 13,947 bp inverted repeat (Figure 5D).

**Figure 5.**
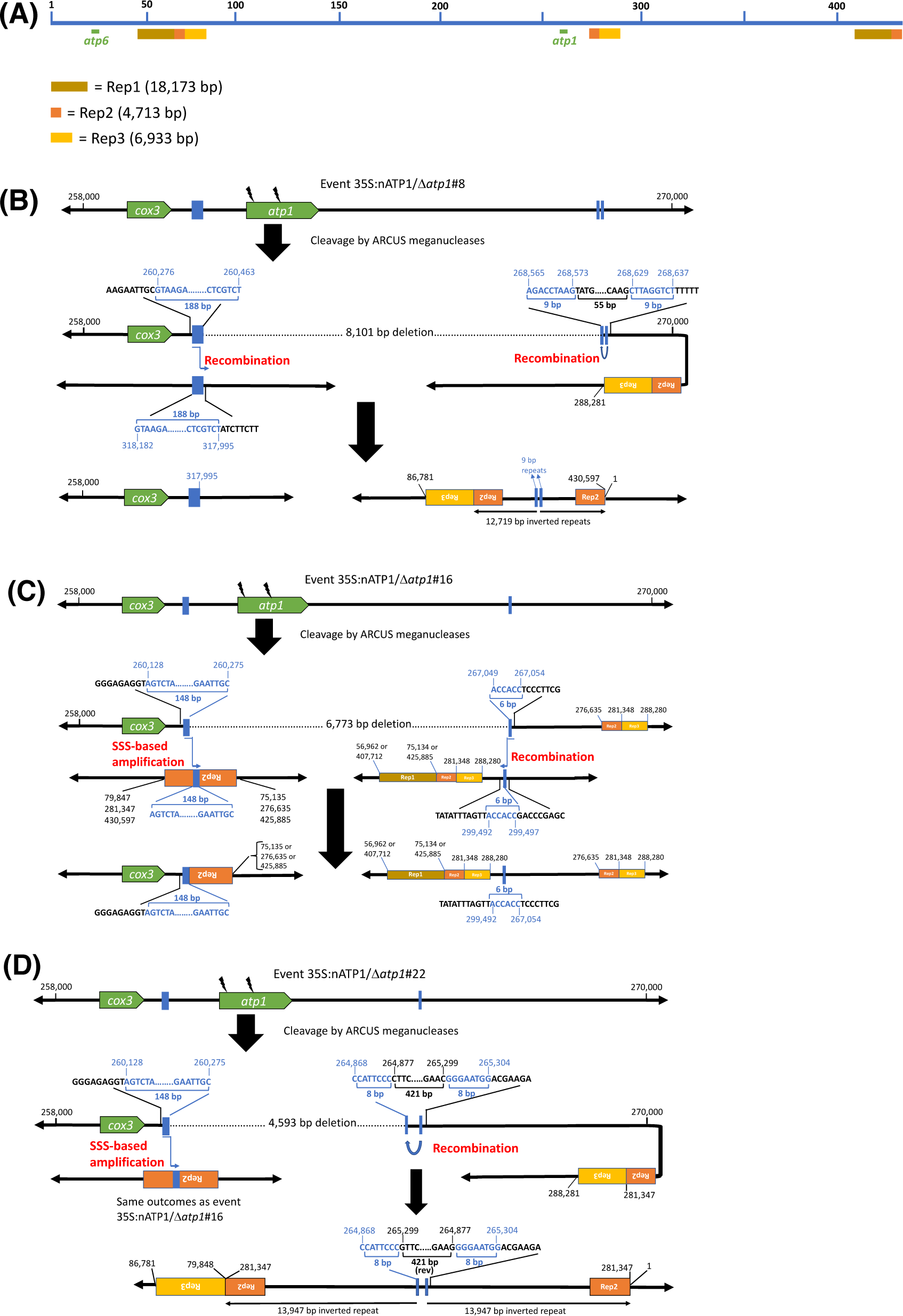
Predicted recombination and SSS events leading to the elimination of *atp1* in 35S:nATP1/Δ*atp1* plants. **(A)** Linear representation of the tobacco mitochondrial reference genome BA000042 showing the locations of *atp1*, *atp6* and the three long direct repeats Rep1, Rep2 and Rep3. Junctions 5’ and 3’ of *atp1* derived via recombination or SSS are shown for T0 events 35S:nATP1/Δ*atp1*#8 **(B)**, 35S:nATP1/Δ*atp1*#16 **(C)**, and 35S:nATP1/Δ*atp1*#22 **(D)**. Lightning bolts depict mitoARCUS cleavage sites. Blue boxes and nucleotides indicate the homologous regions across which recombination is predicted to have occurred in response to mitoARCUS cleavage of *atp1*, or where pre-existing species were amplified by SSS. Nucleotide positions are in accordance to BA000042. Predictions of more than one possibility adjacent to one the major repeats shown in **(C)** and **(D)** are supported by PacBio sequencing-derived contigs and/or PCR and DNA sequence analysis.

PCR and DNA sequence analysis was used to validate the predicted recombination events in the three Δ*atp1* lines. For line 35S:nATP1/Δ*atp1*#8, PCR primers were designed that specifically amplified both the 5’ and 3’ predicted recombination events, and DNA sequence analysis of the amplification products validated the configuration of the sequences predicted by the contigs (Figure 6A). Similarly, the predicted 3’ recombination junctions for events 35S:nATP1/Δ*atp1*#16 and #22 were confirmed by PCR and DNA sequence analysis (Figure 6B). As mentioned above, the predicted 5’ recombination event(s) shared by lines 35S:nATP1/Δ*atp1*#16 and #22 could either extend from the copy of Rep2 that lies ∼15 kb downstream of *atp1*, or from either of the two distal locations where Rep2 abuts directly with Rep1 (Figure 5A). Interestingly, when primers were designed to test each of those options, amplification and sequence validation were not restricted to lines 35S:nATP1/Δ*atp1*#16 and #22, but were also observed in DNAs isolated from 35S:nATP1/Δ*atp1*#8 and WT K326 (Figure 6C).

**Figure 6.**
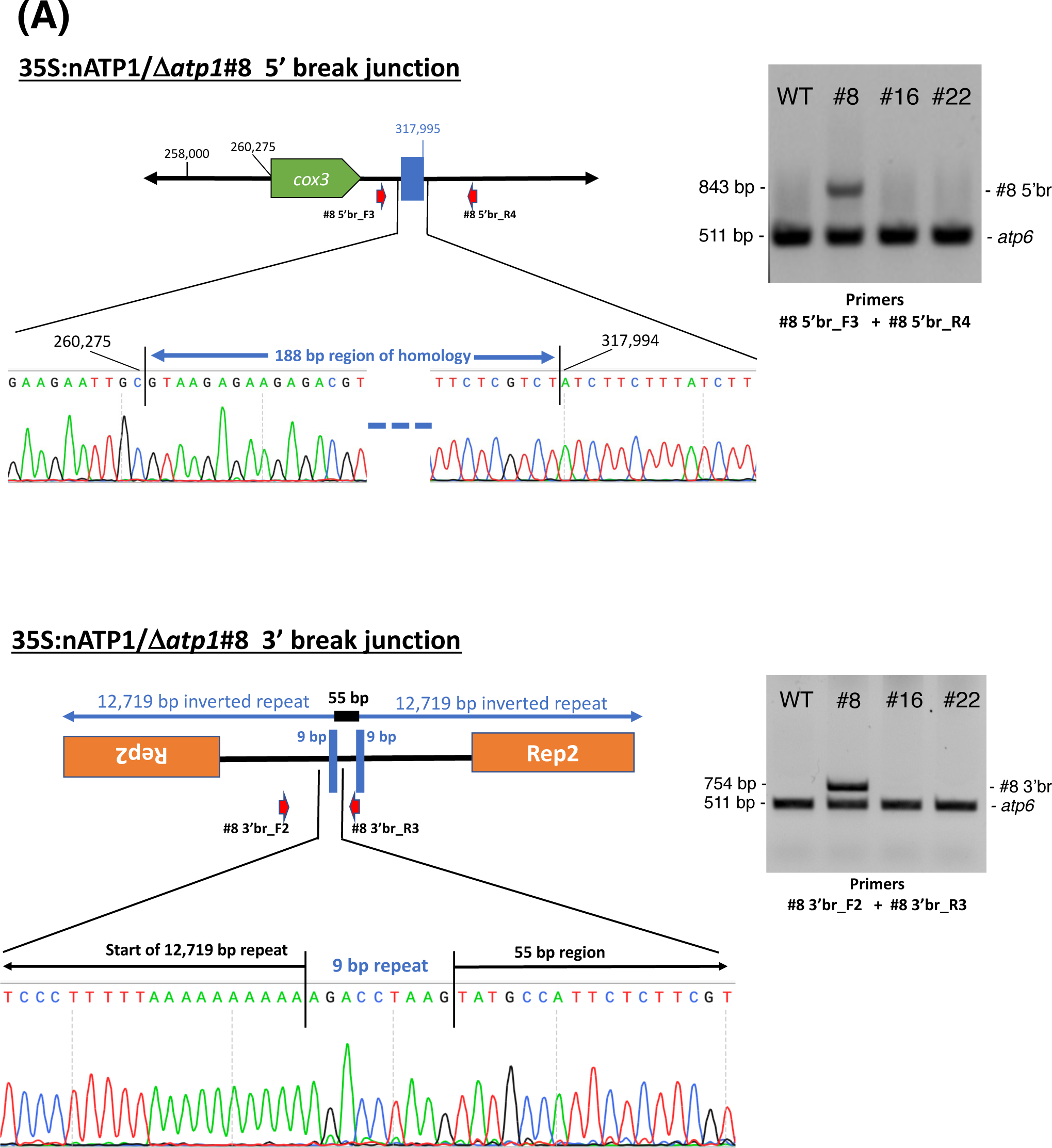
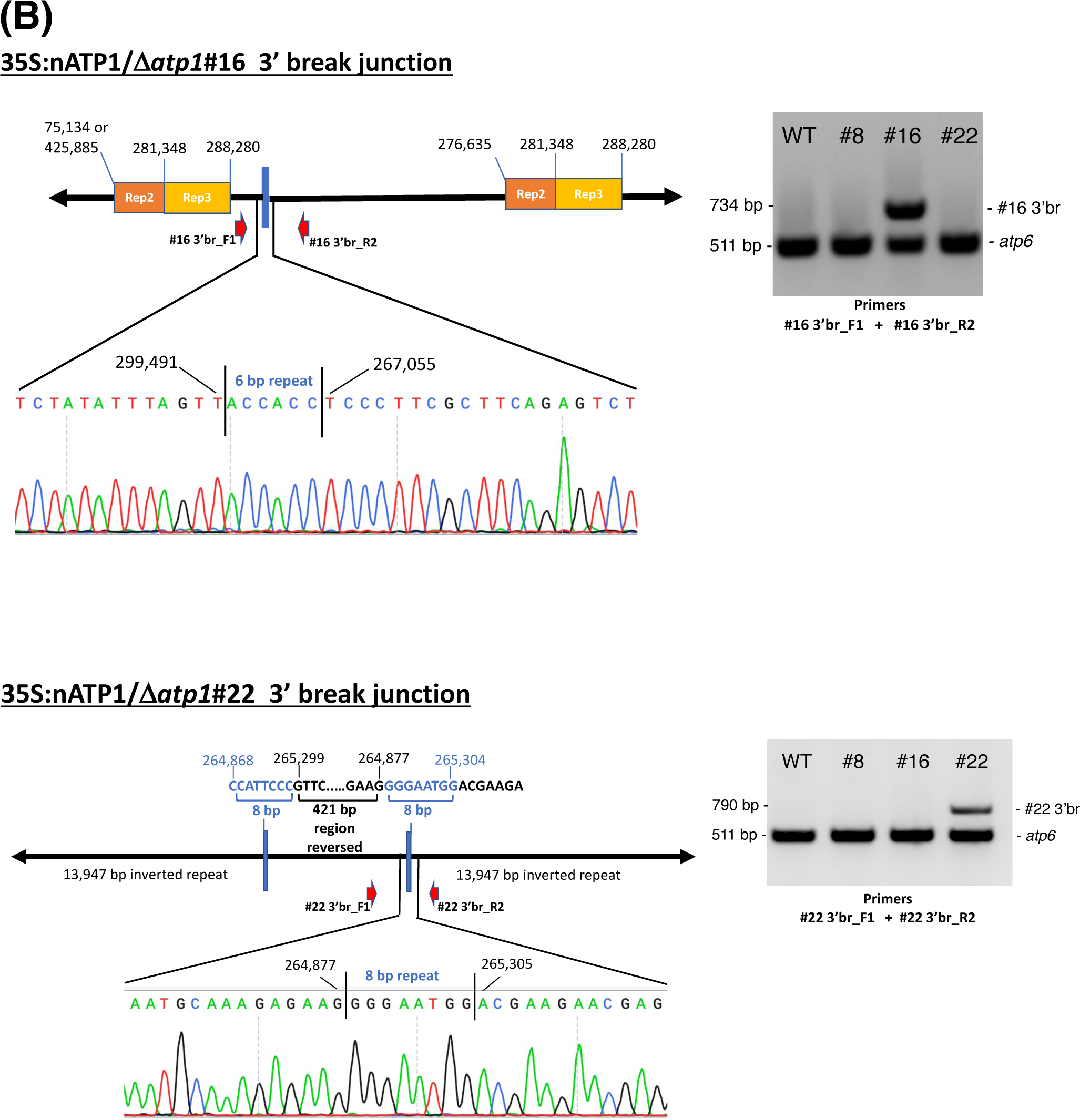
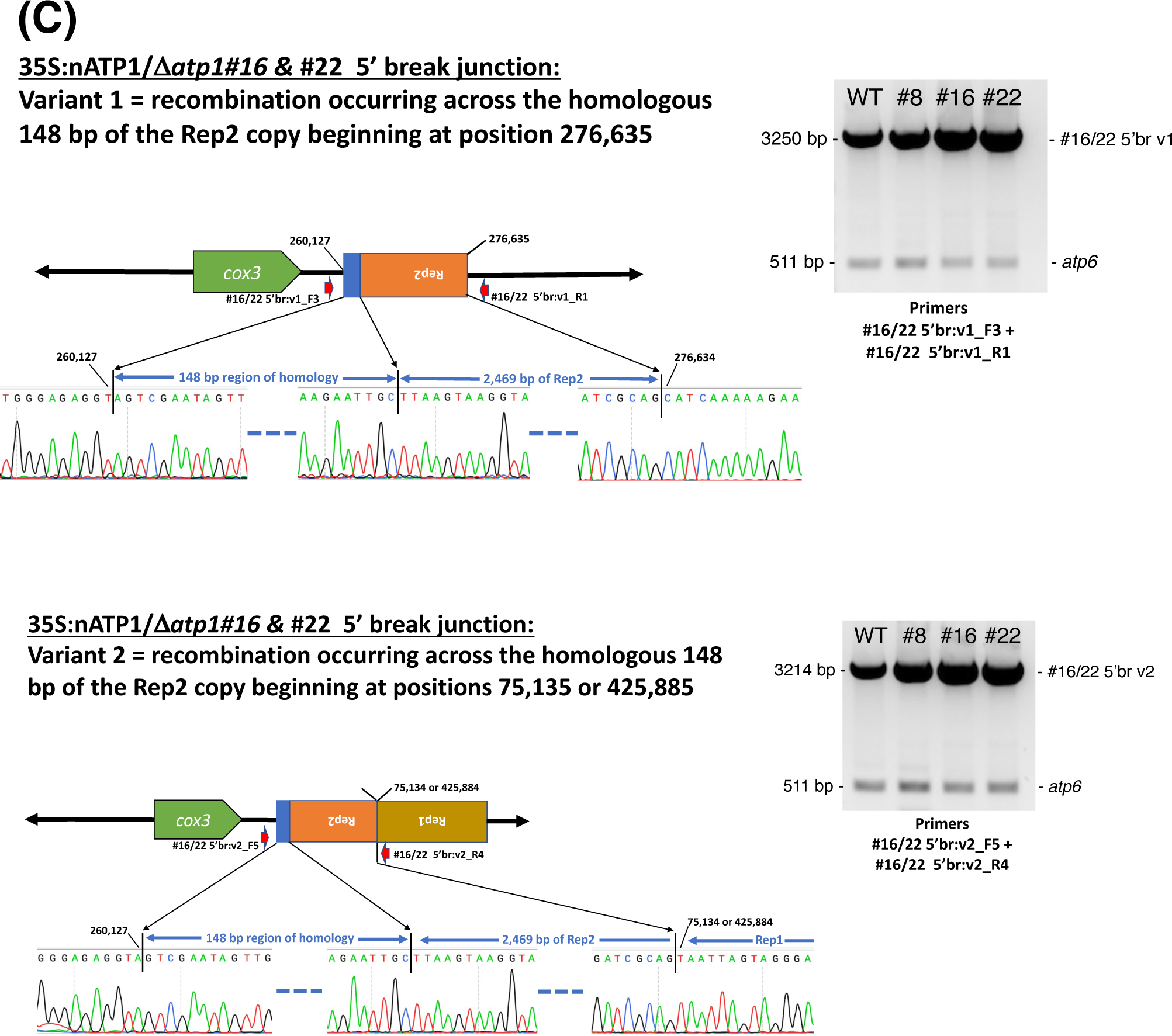
PCR and DNA sequence validation of predicted recombination events in plants possessing Δ*atp1* mutations. **(A)** Recombination junctions 5’ and 3’ of *atp1* in 35S:nATP1/Δ*atp1*#8. **(B)** Recombination junctions 3’ of *atp1* in 35S:nATP1/Δ*atp1*#16 and 35S:nATP1/Δ*atp1*#22. (C) Junctions of pre-existing subgenomic species created by recombination across a 148 bp fragment between *cox3* and *atp1* and the interior of Rep2. Numbering is in accordance with reference genome BA000042. Portions of the DNA chromatograms derived from sequence analysis of the unique PCR products shown in the adjacent gels that validate the recombination junctions predicted in PacBio-derived contigs are shown. For the PCR experiments shown in **(C)**, the *atp6-*specific primers were diluted 1:10 (1 μM final concentration) to prevent the smaller PCR product from outcompeting the amplification of the much larger alternative PCR product.

In their characterization of the tobacco mitochondrial genome, Sugiyama *et al*. (2005) predicted the organization of the genome to be multipartite, comprised of the master circle, as well as subgenomic species generated via recombination across each of the three large repeats (Rep1, Rep2, and Rep3). In addition, they speculated that it was also possible that more subgenomic species may exist as a result of recombination across any of the numerous smaller repeats scattered throughout the genome. Our results suggest that there is indeed a subgenomic species in tobacco mitochondria that is generated through recombination between the 148 bp sequence between *cox3* and *atp1*, and the homologous sequences in reverse orientation found in Rep 2. Our observations suggest that the preservation of essential mitochondrial genes (such as *cox3*) in the region immediately upstream of *atp1* in lines 35S:nATP1/Δ*atp1*#16 and #22 may have not occurred through recombination across DNAs broken by the mitoARCUS enzymes, but rather through the phenomenon known as substoichiometric shifting (SSS), where subgenomic mitochondrial DNA species in low representation can become amplified (Arrieta-Montiel and Mackenzie 2010).

We speculated that the presence of naturally occurring subgenomic species with the recombination junctions predicted 5’ of *atp1* as shown in Figures 5C and 5D could be further tested through examination of the 35S:nATP1/Δ*atp1*#8 PacBio sequence database. Because the 148 bp direct repeat involved in recombination within Rep2 repeats lies immediately upstream of the 188 bp repeat used to create the unique 5’ break junction characterized in line 35S:nATP1/Δ*atp1*#8, there should be PacBio reads in this background that could confirm the presence of these subgenomic species. BLASTN analysis of the PacBio database revealed 11 reads consistent with this hypothesis (Data S6). In nine cases, the sequences at the end of Rep2 were followed by those beginning at position 276,635 (i.e., recombination occurred through the central Rep2); for the other two reads, the sequences at the end of Rep2 were followed by Rep1 (positions 75,135 or 425,885).

### Analysis of 35S:nATP1 X 35S:nATP1/Δ*atp1*#8 progeny

Although our original attempts to germinate seeds obtained from T_0_ 35S:nATP1/Δ*atp1* individuals fertilized with WT or 35S:nATP1 pollen failed to produce a single plant, as tissue-culture-derived clones of the T_0_ plants became available, even more crosses were made using 35S:nATP1 pollen. Seed quantities ranging from 200 – 500 mg were obtained from crosses to the following seven individual 35S:nATP1/Δ*atp1* lines: 35S:nATP1/Δ*atp1*#1, #3, #8, #11, #21, #22, and #23. Consistent with our previous observations, no germination was observed from seeds obtained from six of the seven crossing combinations. However, a total of six plants emerged from an estimated 50,000 seeds planted from the cross 35S:nATP1 X 35S:nATP1/Δ*atp1*#8 (Figure 7). To confirm that the plants were not contaminants, each individual was genotyped using *atp1* primers. All six plants showed the diagnostic pattern of diminished amplification of *atp1-*like sequences in comparison to the co-amplified *atp6* control (Figure 7B). Using primers directed against sequences encoding the ARCUS meganucleases and the *nptII* selectable marker gene, it appeared that the genome editing construct was lost through segregation in two of the six F_1_ plants, and retained in the other four (Figure 7C and 7D). All six F_1_ generation individuals were also assayed using primers specific for the 5’ and 3’ recombination junctions defined in the original 35S:nATP1/Δ*atp1*#8 parent. Interestingly, only three of the six plants showed amplification of the unique 5’ recombination junction, and none of the plants showed the diagnostic amplification product originally observed at the 3’ recombination junction (Figure 7E and 7F). Because subgenomic species created by recombination across the 148 bp region between *cox3* and *atp1* and Rep2 were shown to exist in 35S:nATP1/Δ*atp1*#8 mitochondria, as described above, we speculated that SSS may have resulted in the replacement of species created through recombination across the adjacent 188 bp region with the pre-existing species generated via recombination across the 148 bp repeat shared with Rep2. Semi-quantitative PCR analysis supported this conjecture, as the three F_1_ plants that were PCR positive for the 35S:nATP1/Δ*atp1*#8-specific 5’ recombination product amplified the two subgenomic variants shown in Figure 6c weakly, while those that failed to amplify the former, strongly amplified the latter (compare Figure 7E with Figure S11).

**Figure 7.**
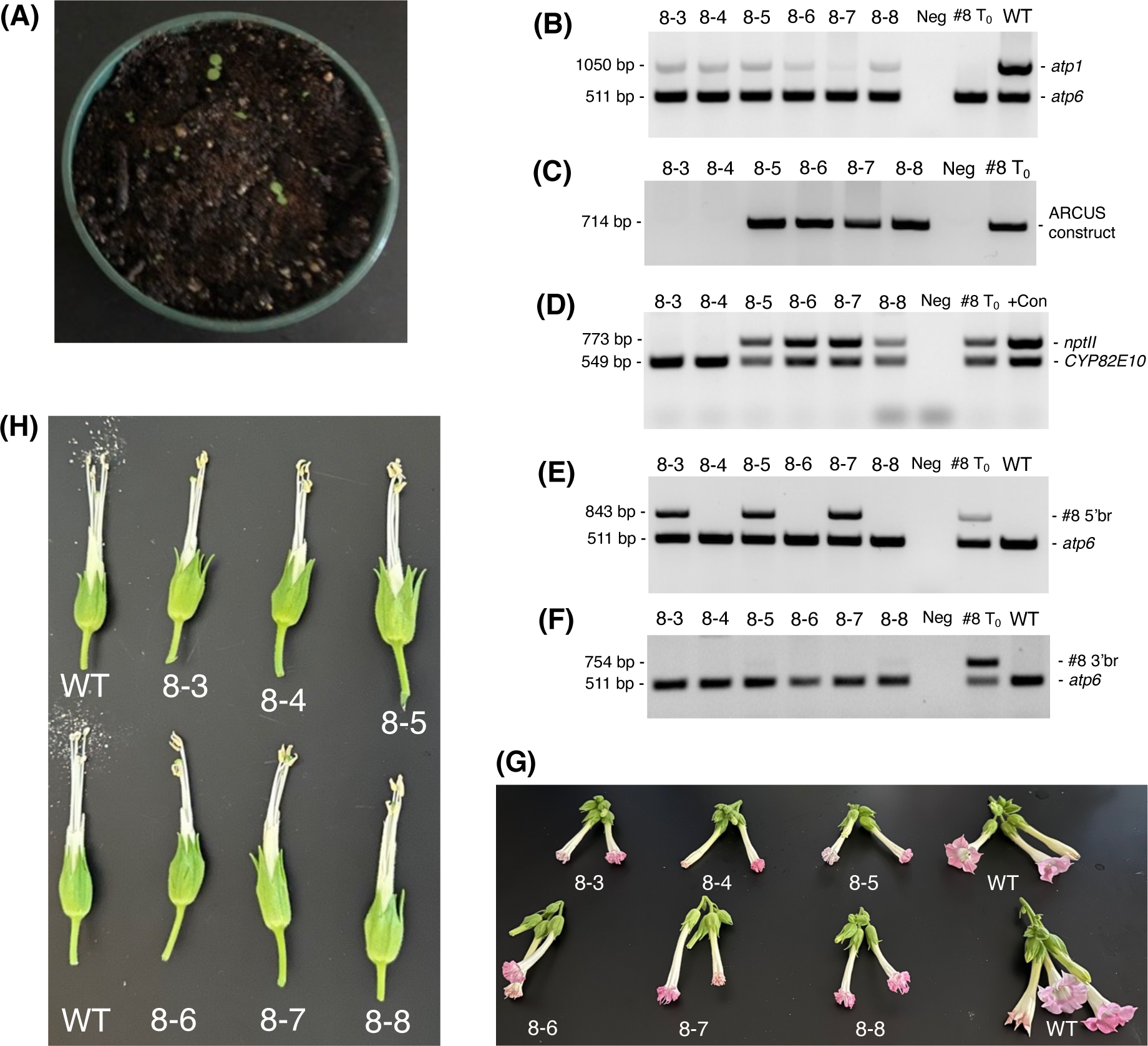
Analysis of 35S:nATP1 X 35S:nATP1/Δ*atp1*#8 F1 progeny. **(A)** Germination of ∼500 mg of 35S:nATP1 X 35S:nATP1/Δ*atp1*#8 seed on potting soil. PCR analysis of six 35S:nATP1 X 35S:nATP1/Δ*atp1*#8 progeny, designated 8-3 through 8-8, using primers designed to amplify: *atp1* **(B)**, the *atp1*-targeting ARCUS transgene **(C)**, the *nptII* selectable marker gene and a control endogenous tobacco gene (*CYP82E10*) **(D)**, and the 5’ break junction **(E)** and the 3’ break junction **(F)** unique to T0 event 35S:nATP1/Δ*atp1*#8. Flowers and anthers of F1 generation Δ*atp1*#8 plants are shown in **(G)** and **(H)**, respectively. The specific primers used in **(B)** through **(F)** are found in Table S1. WT, wild type K326; +Con, unrelated transgenic plant used as *nptII* positive control; Neg, negative control with no DNA template included in the reaction.

When grown to maturity, five of the six 35S:nATP1 X 35S:nATP1/Δ*atp1*#8 F_1_ progeny looked phenotypically normal through the onset of flowering, and one individual displayed a reduced growth habit (Figure S12). Upon flowering, all six F_1_ individuals produced flowers that did not fully expand, and developed mishappen anthers that failed to shed pollen (Figure 7G and 7H).

## DISCUSSION

The endosymbiosis theory is one of the cornerstones of evolutionary biology. In its simplest form, this theory posits that mitochondria evolved from an α-proteobacterium-like ancestor (Timmis *et al*. 2004). After becoming engulfed in a separate host species (the nature of which remains a topic of considerable debate; Martin *et al*. 2015), the great majority of the genes of the organelle progenitor were transferred to the nucleus, with the genomes of modern-day plant mitochondria retaining only a small number of genes (mostly encoding tRNAs, rRNAs, ribosomal proteins, and subunits of enzyme complexes involved in oxidative phosphorylation; Kubo and Newton 2008). Despite the importance of the endosymbiosis theory in our understanding of the evolution of eukaryotic species, to our knowledge this phenomenon had yet to be recapitulated in the lab in a plant species. Using editing tools that target the mitochondrial genome, we have demonstrated the ability to continue the transfer of essential genes from the mitochondria to the nucleus, by repurposing the *atp1* gene to function in the nucleus followed by the complete excision of the endogenous mitochondrial version. These capabilities open new avenues for investigating mitochondrial gene function and genome evolution.

Although plant mitochondrial genomes are commonly depicted as single master circles, *in planta* their structure is multipartite, comprised of populations of circular, linear and even branched species (Chen *et al*. 2017). The concept of subgenomic mitochondrial DNA species becoming amplified or reduced through SSS was initially proposed by Small *et al*. (1987, 1989). In common bean, a CMS-associated subgenomic mitochondrial species estimated to exist at a frequency of less than one copy per 100 cells in some *Phaseolus* species was shown to be amplified 1000-to 2000-fold in others (Arrieta-Montiel *et al*. 2001). Prolonged exposure of plant cells to tissue culture, and downregulation of nuclear genes that function to suppress ectopic recombination of mitochondrial DNAs are two factors known to enhance the frequency of SSS events (Arrieta-Montiel *et al*. 2010). Our results suggest that in addition to spontaneous recombination across regions of microhomology, SSS is another mechanism by which plants restructure their genomes in response to attack by DNA break-inducing endonucleases introduced into the mitochondria.

Analyses of the six 35S:nATP1 X 35S:nATP1/Δ*atp1*#8 F_1_ individuals that were recovered suggested that the recombination events that emerge in the T_0_ generation may not necessarily be the most favored over time. All six of the F_1_ plants showed little to no amplification using the PCR primers that were diagnostic for the recombination event 3’ of *atp1* in the original T_0_ plant. Large inverted repeats in close proximity are typically not found in plant mitochondrial genomes. Thus, it is possible that the 12,719 bp inverted repeat generated in the T_0_ plant (separated by only 55 bp) was not stable through meiotic reproduction. Furthermore, PCR analysis using the diagnostic primers 5’ of *atp1* showed that the band observed in the original T_0_ event was lost in three of the six F_1_ progeny, a loss that appeared to be compensated by SSS-mediated amplification of the two subgenomic variants characterized in 35S:nATP1/Δ*atp1*#16 and #22 (Figure 6C).

For an allotopically expressed mitochondrial gene to completely substitute for the endogenous gene it is replacing, it must be expressed within the nucleus at sufficient levels and in all cell types where the mitochondrial counterpart functions. For mitochondrial genes required for ATP production, this is arguably every living cell. Because our goal was to explore new mechanisms for producing the CMS trait, the ideal promoter would be one that expresses adequately in all cell types except those required for pollen production. Our results support the conclusion that the widely used CaMV 35S promoter is minimally expressed in male reproductive tissues, as all 35S:nATP1/Δ*atp1* plants recovered produced deformed anthers and showed no evidence of pollen production.

In addition to displaying CMS phenotypes, when crosses were made to 35S:nATP1/Δ*atp1* plants they failed to produce viable seed (with the exception of the six F_1_ plants described above). Early expression studies using CaMV 35S promoter:β-glucuronidase (GUS) reporter constructs detected GUS staining throughout the embryo and endosperm in the seeds of transgenic tobacco (Benfey *et al*. 1989) and rice (Terada and Shimamoto, 1990). Both studies, however, used mature seeds as the explant for analysis. When a detailed developmental survey was conducted in cotton plants expressing a green fluorescent protein (GFP) driven by a CaMV 35S promoter, intense GFP fluorescence was observed in both the embryo and endosperm during mid through late seed development, but was undetectable during early seed development (Sunilkumar *et al*. 2002). Our results suggest that tobacco is likely similar to cotton in this respect, and that the 35S promoter is not expressed at a high enough level during early seed development for the 35S:nATP1 construct to compensate for the missing *atp1* gene, leading to aborted seed development. However, the fact that we were able to recover six plants from hundreds of thousands of seeds obtained from seven distinct 35S:nATP1 X 35S:nATP1/Δ*atp1* lines, suggests that on rare occasion it is possible for the 35S:nATP1 construct to generate enough Atp1 to produce viable seeds.

In principle, the strategy described herein to develop a novel CMS trait in tobacco should be broadly applicable across plant species capable of being genetically transformed, and using any essential mitochondrial (or even chloroplast) gene that can be effectively repurposed to function as a nuclear gene. Unless a promoter that is expressed at sufficient levels in all cell types except those required for male reproduction can be identified, however, additional promoter engineering and/or expression strategies will be needed for this CMS trait to be put into practice. Though multiple strategies can be envisioned, for the specific proof-of-concept example described here, one simple approach would be to couple the 35S:nATP construct with another cassette where the nATP1 transgene is placed under the control of a seed-specific promoter that expresses in early stages of seed development.

To create a CMS-based three line system commonly used for hybrid seed production in crops where the seed is the harvested commodity, both maintainer and restorer lines would be required (Chen and Liu, 2014). In our system, the maintainer line would simply be a line in the same genetic background as the CMS line (containing the same complement of allotopic gene constructs), but in a normal cytoplasm. A restorer line would be created by transforming a different inbred with the repurposed mitochondrial gene (nATP1 in our example) under the transcriptional control of a promoter that is highly expressed in all cell types, including those required for male reproduction. Candidate promoters for driving strong constitutive expression throughout all tissues and stages of development would include those associated with *Actin* or *Ubiquitin* genes (Potenza *et al*. 2004; Hernandez-Garcia and Finer 2014). A diagram depicting the proposed three-line system based on the creation of CMS traits using allotopic expression and mitochondrial genome editing is shown in Figure 8. It should be noted that although the proof-of-concept demonstration described here, and the model presented in Figure 8, focus on inactivating an essential gene located in the mitochondria, in principle one should also obtain similar outcomes through the allotopic expression and knockout of an essential chloroplast gene.

**Figure 8.**
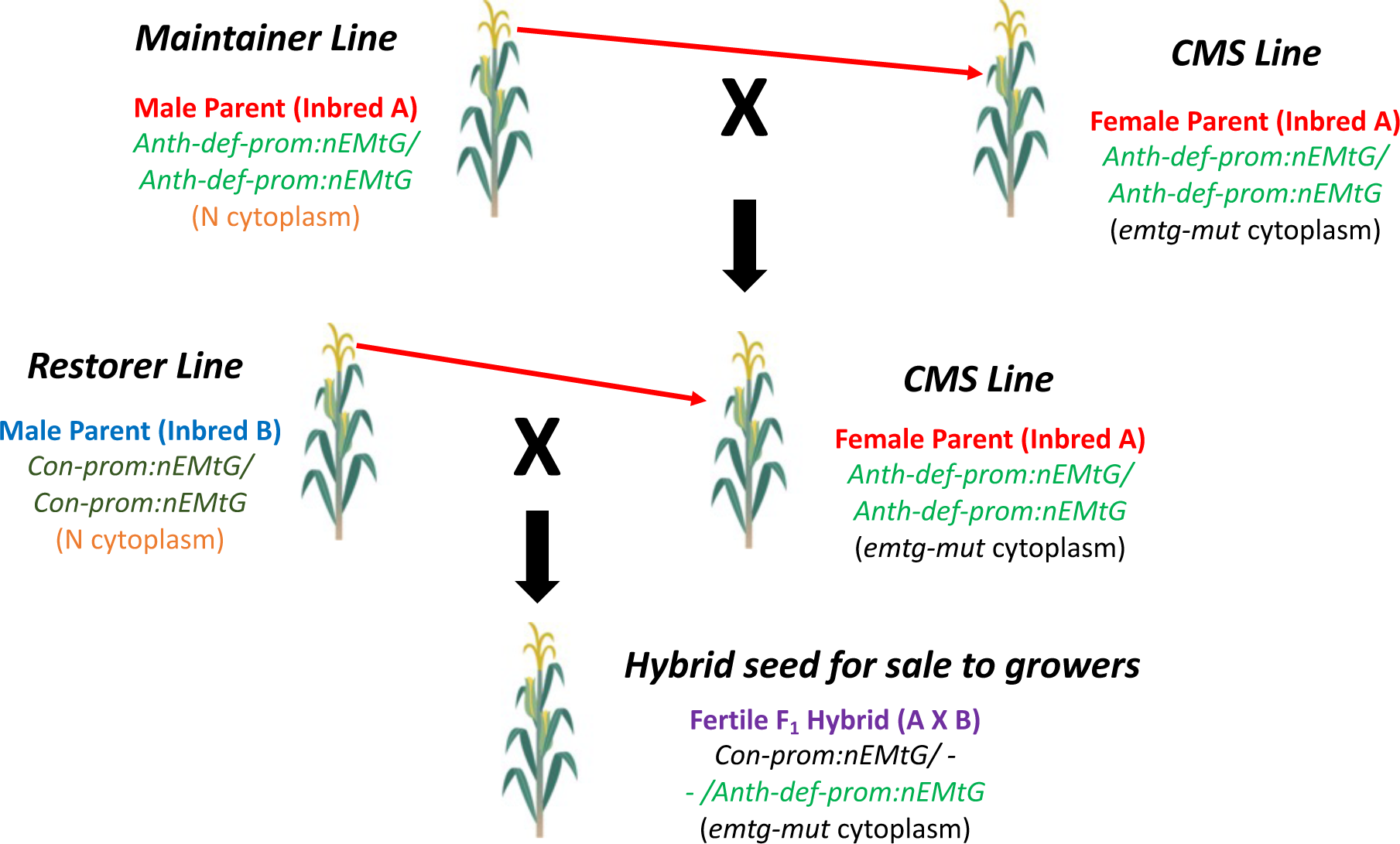
Proposed CMS-based hybrid seed production system based on allotopic expression combined with mitochondrial genome editing. The CMS inbred is propagated by crossing to the maintainer line as the pollen parent; hybrid seed is produced by crossing the CMS line with the restorer line. Fertility in the field is restored by the Con-prom:nEMtG construct. Anth-def-prom, promoter expressed in all tissues except anthers; Con-prom, promoter expressed in all plant tissues; nEMtG, essential mitochondrial gene repurposed to function in the nucleus; emtg-mut, essential mitochondrial gene rendered nonfunctional through genome editing. Maize is depicted as the example crop species.

Capsule development appeared normal when 35S:nATP1/Δ*atp1* CMS plants were fertilized with either 35S:nATP1 or WT pollen, suggesting that a normal program of ovary development was initiated post-fertilization. The seed produced within these capsules, however, were smaller, lighter in color and weight, and appeared to be largely devoid of cellular content. This phenotype displays the hallmarks of stenospermocarpy, a form of seedless fruit production.

Stenospermocarpy differs from parthenocarpy, another mechanism through which a seedless phenotype may be manifest, in that seedless fruits arise in the absence of fertilization in the latter, whereas fruit development requires fertilization for the former to occur (Joldersma and Liu 2018). Bananas, citrus, cucumbers and pineapples are examples of commodities where seedlessness can be achieved through parthenocarpy; seedless watermelons and grapes are popular examples whereby seedlessness is produced via stenospermocarpy (www.canr.msu.edu/news/seedless-fruit-is-not-something-new). Although fruits developed through stenospermocarpy are not literally without seeds, the abortive seed remnants are small and soft enough that consumers do not mind them. Due to the popularity of seedless fruit phenotypes, substantial efforts have been devoted toward understanding the molecular mechanisms underlying the trait. The great majority of genes whose function can be altered to yield a seedless phenotype affect hormone production, histone methylation, or signaling pathways (Joldersma and Liu 2018). Thus, it is not surprising that the seedless trait is often associated with pleiotropic phenotypes, such as reduced fruit size, mishappened leaves or fruit, and reduced yield when grown in the field (Gorguet *et al*. 2005; Mazzucato *et al*. 2015). Should the seedless trait observed in fertilized 35S:nATP1/Δ*atp1* tobacco plants be similarly observed when the system is transferred to a fruiting species such as tomato (a close relative of tobacco), this could represent a means for generating seedless phenotypes in a manner that is mechanistically distinct from other strategies described to date. From a commercial perspective, seedless phenotypes are most easily managed in species that can be readily propagated vegetatively through micropropagation or cuttings. Should the system described here prove viable for seedless fruit production in other crops, it would be advantageous to couple the 35S:nATP1 cassette with another in which the nATP1 construct is under the transcriptional control of a promoter that expresses in male reproductive tissue, and not seeds, as this would eliminate the need for cross-pollination to stimulate ovary/fruit development.

Finally, another potential application of the mitochondrial genome editing technology described here could be for the purpose of transgene containment (see Clark and Maselko, 2020 for a recent review). In cases where there is a need to minimize the possibility of a transgene of interest from being unintentionally propagated in the wild or introduced into a cross-compatible nontransgenic crop, arguably the ultimate form of containment would be to introduce the transgene in a background where transmission through both pollen and seed is severely inhibited. Should a transgene(s) responsible for the production of a high value pharmaceutical or industrial enzyme, for example, be introduced into a 35S:nATP1/Δ*atp1* tobacco background, there would be minimal risk of transgene escape from the vegetatively propagated clonal progeny that would be farmed for the high value product. Furthermore, should the CaMV 35S promoter show a generalized pattern of negligible expression in both male reproductive tissues and developing seeds across a wide range of plant species, the transferal of this system for the purpose of transgene containment could have broad application.

## MATERIALS AND METHODS

### MitoARCUS design

A combination of directed evolution and in silico predictions were used by Precision BioSciences to re-engineer *I-CreI* (including a peptide linker to join the two dimers as a monomer) to bind and cleave 22bp sequences specific to *atp1* as previously described (Smith *et al*. 2012; Smith and Jantz, 2013). Initial confirmation of mitoARCUS specificity was conducted using a split-GFP assay in CHO cells as described in Zekonyte *et al*. (2021).

### Vector construction, tobacco transformation, and molecular characterization

The repurposed *atp1* construct (nATP1), codon optimized for expression as a tobacco nuclear gene and including the sequences encoding the transit peptide and 3’-UTR of *ATP2* (Figure S1), was custom synthesized by GenScript (Piscataway, NJ). nATP1 was cloned downstream of an enhanced CaMV 35S promoter and introduced into plant expression vector pCAMBIA1300 (Abcam) that contains an *hptII* selectable marker gene. The *atp1*-targeting vector SP3379 contains two mitoARCUS cassettes: ATP-5/6, which was cloned downstream of a chimeric *Figwort Mosaic Virus – Peanut Chlorotic Streak Caulimovirus* promoter (Acharya *et al*. 2014) and upstream of the termination sequences of the soybean *Ubiquitin3* gene; and ATP-7/8, under the transcriptional control of an enhanced CaMV 35S promoter and termination sequences mediated by the soybean *MYB2* gene (Figure S3). Transport of the ARCUS enzymes into the mitochondria was facilitated by a transit peptide designated M20, derived from a putative Arabidopsis HNH endonuclease (GenBank accession KAG6466785). SP3379 contains an *nptII* selectable marker gene.

Tobacco haploids were generated by crossing variety K326 with pollen from *N. africana* (Burk *et al*. 1979). Vector pCAMBIA1300-35S:nATP1 was introduced into *Agrobacterium tumefaciens* strain LBA4404 and transformed into haploid K326 leaf discs using standard protocols (Horsch *et al*. 1985) and hygromycin selection. Several independent T_0_ haploids were screened using semi-quantitative PCR to identify individuals accumulating the highest levels of nATP1 transcripts. Selected haploids were chromosome doubled using the midvein culturing technique (Kasperbauer and Collins 1972) to fix the transgene to homozygosity. Several doubled haploid individuals were again screened by semi-quantitative PCR to confirm high transgene expression, and plant 35S:nATP1#25 was chosen for further experiments.

MitoARCUS vector SP3379 was transformed into 35S:nATP1#25 using the same *Agrobacterium*-based transformation protocol, but with kanamycin selection. As an EV control, a K326 doubled haploid plant transformed with the pCAMBIA1300 vector alone was also used as a host for transformation with SP3379.

For PCR analyses, total cellular DNA was isolated from tobacco leaf tissue using the MP Biomedicals FastDNA Kit. Primer sequences and reaction conditions for all PCR experiments are listed in Table S1. DNA sequence analysis of PCR amplification products was conducted by the NCSU Genomic Sciences Laboratory (https://research.ncsu.edu/gsl).

### Imaging of anthers, pollen and seeds

Tobacco pollen was stained within intact anthers as previously described (Peterson *et al*. 2010) and imaged using a Labomed CxL microscope with an AmScope MU1000-HS digital camera. Toluidine Blue-O (TBO; Sigma-Aldrich) staining was performed on developmentally staged anthers or ovaries as previously described (Strable *et al*. 2020). Briefly, TBO was dissolved in 1% sodium borate (w/v) to make a 0.5% TBO staining solution (w/v). Slide-adhered microtome sections (10 µm) were deparaffinized in Histo-Clear (National Diagnostics). Slides were hydrated through a graded ethanol series, 1 min each (100%, 100, 95, 95, 70, 50, distilled water) and stained in 0.5% TBO staining solution for 1 min. Slides were then dehydrated through a graded ethanol series, 30 sec each (distilled water, 50%, 70, 95, 95, 100, 100) and Histo-Clear (three times, 1 min each). Slides were dried briefly and mounted with Permount (Fisher) and a cover slip. A Leica DM4B microscope equipped with a DMC6200 digital camera was used to capture images. Images of intact anthers and stigmas, and tobacco seeds germinated on solid MS media were taken using a dissecting microscope (Nikon SMZ-U) with an AmScope MU1000-HS digital camera.

### PacBio sequencing and analysis

A scaled-up version of the initial steps of the protocol described by Ahmed and Fu (2015) was used to enrich the preparations for mitochondria prior to DNA exaction. Briefly, thirty grams of leaf tissue from lines 35S:nATP1/Δ*atp1*#8, #16, and #22 were ground on ice using a pre-chilled grinding medium (0.5M sucrose, 1 mM EDTA, 70 mM KH_2_PO_4_, pH 7.5) supplemented with 0.8 % BSA and 0.1% β-mercaptoethanol. After filtering through two layers of Miracloth (Sigma-Aldrich), the homogenate was centrifuged at 1800*g* for 7 min to pellet intact nuclei, chloroplasts, and cellular debris. The supernatant was centrifuged at 17,200*g* for 20 min to pellet a fraction enriched in mitochondria. The pellet was resuspended in grinding media plus 0.8% BSA and the cycle of discarding the pellet of a low-speed spin (1800*g*), resuspension, and retention of the pellet after a high-speed spin (17,200*g*) was repeated two more times. DNA was extracted from the final pellets using the NucleoBond High Molecular Weight DNA Kit according to the manufacturer’s instruction (Macherey-Nagel).

Library preparation and PacBio sequencing was conducted by Eremid Genomic Services (Kannapolis, NC) using the SMRTbell prep kit 3.0 (PacBio), following the manufacturer’s instruction. In brief, for each sample, 10 µg of high molecular weight genomic DNA was fragmented using g-TUBE (Covaris), ligated to a SMRTbell barcoded adapter 3.0 and size selected using AMPure PB beads (PacBio) to remove fragments shorter than 5 kb. The integrity of each library was analyzed on TapeStation (Agilent), pooled in equimolar concentration, and loaded by diffusion at 350 pM into one SMRTcell-8M. Sequencing was performed on the Sequel-IIe system HiFi-CCS mode (PacBio).

HiFi reads were mapped to *N. tabacum* mitochondrial reference genome BA000042 using Minimap2 (version 2.24-r1122) with the flags-ax map-hifi--sam-hit-only (Li 2018, 2021) to estimate the number of reads of potential mitochondrial origin. The alignment results were summarized with samtools (v 1.12) to calculate the percentage of reads that aligned to BA000042 (Danecek *et al*. 2021). Read coverage was analyzed using the Integrative Genomics Viewer (IGV, v 2.12.0) (Robinson *et al*. 2011) and the R package karyoploteR (v 1.20.3) (Gel and Serra 2017). To help reduce the number of NUMTs and chloroplast sequences included in analyses of the mapped reads, reads that were more than 33% clipped were removed. These reads were identified using the SamJdk program from JVarkit (Lindenbaum 2015).

For each of the three Δ*atp1* genotypes, the HiFi reads that mapped to the reference genome were assembled into contigs using hifiasm with the flag −l 0 (version 0.16.1-r375)(Cheng *et al*. 2021; 2022). Each assembly was then aligned to BA000042 using nucmer (v mummer-4.0.0rc1) with the -c 250 flag (Delcher *et al*. 2002; Kurtz *et al*. 2004) to identify potential mitochondrial contigs. From this set, contigs that shared greater than 99.99% identity with the reference (based on BLASTN) were classified as representing the mitochondrial genome; contigs whose homology to BA000042 was less than 98% were classified as NUMTs. In six cases, a contig shared greater than 99.99% sequence identity to the reference genome over a region spanning more than 100 kb, but was connected to sequences at the 3’ or 5’ end that were clearly derived from NUMTs. These contigs were hand curated to trim the non-mitochondrial sequences. BLASTN alignments of genuine mitochondrial contigs against BA000042 were conducted to identify the portions of the genome that were missing in the Δ*atp1* mutants, as well as define the nature of the specific junctions 5’ and 3’ of *atp1* where the mutant mitochondrial genomes differed from the reference. To visualize the alignments between mitochondrial contigs and BA000042, circular plots were generated with Circos v 0.69-6 (Krzywinski *et al*. 2009) using nucmer with the -c 250 flag, followed by delta-filter -q (v mummer-4.0.0rc1) to obtain the best hit alignments.

## Acknowledgements

This work was funded in part by a grant from Elo Life Systems to R.E.D. (NCSU grant 2019-1417). The authors are greatly appreciative of the NCSU Plant Breeding Consortium for providing the support for the bioinformatic analyses. We are grateful to Kevin Stroup for the tissue culture and maintenance of T_0_ 35S:nATP1/Δ*atp1* plants, and Eli Hornstein for assistance with the Labomed CxL microscopy and imaging. We also thank Ramsey Lewis for providing the K326 haploid materials used for 35S:nATP1 transformation.

## Author contributions

R.E.D. conceived and designed the project, analyzed the data and wrote the manuscript. D.S., H.C.G. and W.A.S. conducted most of the experiments. A.N.D. analyzed the PacBio sequence data and wrote the corresponding methods. D.J. and J.J.S. designed the two ARCUS meganucleases. A.M. tested the M20 transit peptide and constructed the SP3379 vector. J.S. and C.K. conducted the histological sectioning and staining, and J.S. wrote the corresponding methods. All authors read and approved the final manuscript.

## Supporting information

**Figure S1** DNA sequences of the native mitochondrial *atp1* gene and the repurposed 35S:nATP1 construct designed to function as a nuclear gene.

**Figure S2** Semi-quantitative PCR analysis of K326 haploid plants transformed with 35S:nATP1.

**Figure S3** MitoARCUS vector SP3379.

**Figure S4** Transient expression of ARCUS-GFP fusion proteins in tobacco protoplasts.

**Figure S5** Primer combinations designed to amplify *atp1*.

**Figure S6** Examples of mature T_0_ 35S:nATP1/Δ*atp1* plants.

**Figure S7** Examples of normal and 35S:nATP1/Δ*atp1* flowers and anthers.

**Figure S8** Mature capsules (a) and seeds (b) of plant 35S:nATP1/Δ*atp1*#8 fertilized with WT tobacco pollen.

**Figure S9** Plants used for PacBio sequence analysis.

**Figure S10** Circos plots of contigs corresponding to the mitochondrial genomes of 35S:nATP1/Δ*atp1*#8 (a), 35S:nATP1/Δ*atp1*#16 (b), and 35S:nATP1/Δ*atp1*#22 (c) aligned to tobacco mitochondrial reference genome BA000042.

**Figure S11** SSS in 35S:nATP1 X 35S:nATP1/Δ*atp1*#8 F_1_ progeny.

**Figure S12** 35S:nATP1 X 35S:nATP1/Δ*atp1*#8 F_1_ plants at flowering.

**Table S1** PCR primers and reaction conditions used in this study.

**Table S2** PacBio sequence assembly summary.

**Data S1** Single run DNA sequences of independently cloned PCR products of WT K326 DNA amplified with *atp1* primers.

**Data S2** Single run DNA sequences of independently cloned PCR products of plant 35S:nATP1/Δ*atp1*#8 amplified with *atp1* primers.

**Data S3** Single run DNA sequences of independently cloned PCR products of plant 35S:nATP1/Δ*atp1*#16 amplified with *atp1* primers.

**Data S4** *atp1*-like reading frames of tobacco NUMTs found in GenBank (>93% identify over a minimum of 1 kb).

**Data S5** *atp1*-like reading frames of tobacco NUMTs identified in PacBio reads.

**Data S6** PacBio reads from line 35S:nATP1/Δ*atp1*#8 that support the existence of subgenomic species generated through recombination between the 148 bp repeat located between *cox3* and *atp1*, and the interior of Rep2.

## Other data access

Fasta files for the three assemblies and the six trimmed contigs can be found on Dryad (https://datadryad.org/stash/share/ZIYx4FtC9CwFEsdSJCE3c4Cc2wr3RTpytwYVEB5o-cE). The Mt genome contig ids are ptg000007l_trimmed, ptg000008l, ptg000020l_trimmed, ptg000022l_trimmed, and ptg000027l for 35S:nATP1/Δ*atp1*#8; ptg000004l _trimmed, ptg000017l, ptg000024l, and ptg000037l for 35S:nATP1/Δ*atp1*#16; ptg000006l _trimmed, ptg000008l _trimmed, ptg000027l, ptg000032l, and ptg000063l for 35S:nATP1/Δ*atp1*#22

