## Supplementary material for "Cytoplasmic Male Sterility and Abortive Seed Traits Generated through Mitochondrial Genome Editing Coupled with Allotopic Expression of *atp1* in Tobacco": Figure S1

## (A)

**ATG**gaactttctccccgagctgcggaactaacaagtctattagaaagtcgaattagcaacttttacaccaattttca
agtggatgagatcggtcgagtggtctcagttggagatgggattgcacgtgtttatggattgaacgagattcaagctg
gggaaatggttgaatttgccagcggtgtgaaaggaatagccttgaatcttgagaatgagaatgtagggattgttgtc
tttggtagtgatactgctattaaagaaggagatcttgtcaagcgcactggatctattgtggatgttcctgcgggaaa
ggctatgctagggcgtgtggtcgatggcttgggagtacctattgatggaagggggctctaagcgatcacgagcgaa
gacgtgtcgaagtgaaagcccctggtattattgaacgtaaatctgtgcacgagcctatgcaaacagggttaaaagcg
gtagatagcctggttcctataggtcgtggtcaacgagaacttataatcggggaccgacaaactggaaaaactgctat
tgctatcgataccatattaaaccaaaagcaactgaactcaagggccacctctgagagtgagacattgtattgtgtct
atgtagcgattggacagaaacgctcaactgtggcacaattagttcaaattctttcagaagcgaatgctttggaatat
tctattcttgtagcagccaccgcttcggatcctgctcctctacaattttggccccatattctgggtgtgccatggg
ggaatatttccgcgataatggaatgcacgcattaataatctatgatgatcttagtaaacaggcggtagcatatcgac
aaatgtcattattgttacgccgaccaccaggtcgtgaggctttcccaggggatgttttctatttacattcccgtctc
ttagaaagagcggctaaacgatcggaccagacaggcgcaggtagcttgaccgccttacccgtcattgaaacacaggc
tggagacgtatcggcctatattcccaccaatgtgatcccattactgatggacaaatctgtttggaaacagagctct
tttatcgcggaattagacctgcgattaacgtcggcttatctgtcagtcgcgtcgggtctgccgctcagttgaaaact
atgaaacaagtctgcggtagttcaaaactggaattggcacaatatcgcgaagtggccgcccttgctcaatttggctc
agaccttgatgctgcgactcaggcattactcaatagaggtgcaaggctgacagaagtaccgaaacaaccacaatatg
caccactgccaattgaaaaacaaattctagtcatttatgcagctgtcaatggattctgtgatcgaatgccactagac
agaatttctcaatatgagagagccattccaaatagtgtcaaaccagaattactacaatcctttttagaaaaaggtgg
cttaactaacgaaagaaagatggaaccagatacattcttaaaagaaagtgctttagcttttatt**TAA**

## (B)

**gtcgac**gagaggcggtttgcgtattggctagagcagcttgccaacatggtggagcacgacactctcgtctactccaa
gaatatcaaagatacagtctcagaagaccaaagggctattgagacttttcaacaaagggtaatatcgggaaacctcc
tcggattccattgcccagctatctgtcacttcatcaaaaggacagtagaaaaggaaggtggcacctacaaatgccat
cattgcgataaaggaaaggctatcgttcaagatgcctctgccgacagtggtcccaaagatggacccccacccacgag
gagcatcgtggaaaaagaagacgttccaaccacgtcttcaaagcaagtggattgatgtgaacatggtggagcacgac
actctcgtctactccaagaatatcaaagatacagtctcagaagaccaaagggctattgagacttttcaacaaagggt
aatatcgggaaacctcctcggattccattgcccagctatctgtcacttcatcaaaaggacagtagaaaaggaaggtg
gcacctacaaatgccatcattgcgataaaggaaaggctatcgttcaagatgcctctgccgacagtggtcccaaagat
ggacccccacccacgaggagcatcgtggaaaaagaagacgttccaaccacgtcttcaaagcaagtggattgatgtga
tatctccactgacgtaagggatgacgcacaatcccactatccttcgcaagacccttcctctatataaggaagttcat
ttcatttggagaggacacgctgaaatcaccagtctctctctacaaatctatctctctcgagctttcgcagatctgtc
gaacca**ggatcc**act**ATG**gcttctcggaggcttctcacctctctcctccgtcaatcggctcaacgtggcggcggtcc
aatttcccgatccttgggaaactccatccctaaatccgctgcacgcgcctcttcacgcgcgtcccctaagggattcc
tcttaaaccgcgccgtacagtacgctacctctgcagcagcacccgcatctcagccatccatggaactctcaccaagg
gcagcagaacttacttcactcctcgaatctagaatctcaaacttctacacaaactttcaggttgatgaaattggaag
agttgtttcagttggagatggtattgctagggtttatggacttaatgagattcaagctggtgaaatggttgagtttg
cttctggagttaagggtatcgctttgaatcttgaaaatgagaacgttggaatcgttgtttttggttcagatactgct
attaaggagggagatttggttaaaagaacaggttctattgttgatgttccagctggaaaagctatgcttggtagagt
tgttgatggattgggtgttcctattgatggaaggggtgctttgtctgatcatgaaagaaggagagttgaggttaagg
ctccaggaattattgaaagaaaatcagttcatgagcctatgcaaactggtcttaaggctgttgattctttggttcca
attggaaggggtcaaagagaacttattattggagataggcaaactggaaagactgctatcgctatcgatacaatcct
taaccaaaagcaattgaattcaagagctacttctgaatcagagacattgtattgtgtttacgttgctattggacaaa
aaaggtctactgttgctcaacttgttcaaattttgtctgaagctaatgctcttgagtattcaattttggttgctgct
acagcttctgatccagctcctcttcaattttggctccatattcaggatgcgctatgggtgaatactttagagataa
tggtatgcatgctcttattatatatgatgatttgtcaaagcaagctgttgcttacagacaaatgtctcttttgctta
gaagaccaccaggaagggaagcttttcctggagatgttttctatcttcattctaggttgcttgagagagctgctaaa
aggtctgatcaaactggagctggttcacttacagctttgccagttattgagactcaagctggagatgtttcagctta
catccctactaacgttatctctatcacagatggacaaatttgtcttgaaacagagttgttttacaggggtattagac
cagctattaatgttggactttctgtttcaagagttggttcagctgctcaattgaagactatgaaacaagtttgcggt
tctttgaagcttgaattggctcaatatagggaggttgctgcttttgctcaatttggatcagatttggatgctgctac
tcaagctttgcttaatagggggtgctagacttacagaagttttgaagcaacctcaatatgctccacttcctatcgaga
agcaaattcttgttatatatgctgctgttaatggtttctgtgatagaatgccacttgataggatctctcaatacgaa

agagctatcttgaactcagttaagcctgaattgcttcaatcttttcttgagaaaggaggtttgacaaacgagaggaa
gatggagctcgatacttttctcaaggagtccgcactcgcattcatc**TGAagatct***atagattataaacttctgtgac*
*tttcttttcttctctttgccaaaataatttagtttgtgacatcccggatttttttggaggaccaagaggtccagaat*
*tctggttttgttttacatccaatgcgagattatagagacatgcagccaagcctttgttgccagagacccccttttct*
*gttatgtcacataataaaggggtaaatggtgatcttgtatatctgattttcaagtctttttcgagaattttggatt*
*ccctgattatcaaatgcctttctgaaacgctttctttctatatgtggttaacttcacgtccattttatccattctgc*
*cttgaaagcttcaatgtaataggagcagttaatatgctaagcggatacaaatcatatttcttggcgcgacttaattt*
*tattgcagtatgaatacttgtagaaaatatgagtatttgactaagcttataagcggcaaaccagcttataagtcact*
*tttactttacttcatctacgcgtttggtaaaattaaaagtgttaagtctagtgcttgtaagcttttaagtcttaagt*
*tgtcataagttggtcacatctaaatttgaactgcccctgcccccatcttccccaaaagaatctgcattcacgctaag*
*aggttatgcaaactctttagggctaaagaattgtcgtcaaattataactaaagccaaagttctaaaatatgtcttac*
*agagataatgtctagagaataataagatttatatgcagagacaattcctttagattcctttaggatgataataacac*
*acgccagacatacattatagtggaaaaaataatgaacaggaaattacacccaataaccttgaacatatatggtcttg*
*atttttgattctttgtccc***ggtacc**

**Figure S1. DNA sequences of the native mitochondrial *atp1* gene and the repurposed 35S:nATP1 construct designed to function as a nuclear gene. (A)** DNA sequence of the tobacco *atp1* gene. The start and stop codons are in bold, capitalized text. The six 'c' nucleotides shaded grey represent locations where cytidines are converted to uridines in the mature transcript by RNA editing. The 22 bp recognition sites of the ARCUS enzymes ATP-5/6 and ATP-7/8 are shaded green and turquoise, respectively. The shared 4 bp 3' overhangs produced after cleavage are underlined. **(B)** Sequence of construct 35S:nATP1. An enhanced CaMV 35S promoter sequence is shaded yellow. Sequences encoding the tobacco ATP2 transit peptide (plus the first 12 codons of the mature protein) are highlighted in magenta. The 1530 bp sequence in lower case unshaded type represents the custom-synthesized nATP1 reading frame codon optimized for expression in the nucleus of tobacco. The italicized sequences highlighted in green correspond to the region downstream of the tobacco *ATP2* stop codon. Within the green highlighted region, sequences that are underlined can be found in *ATP2* cDNAs, and thus represent the minimal extent of the *ATP2* 3'-UTR. The start and stop codons of the nATP1 open reading frame are indicated in bold, capitalized type. Restriction sights that were engineered to facilitate cloning are shown in lower case, bold type. *ATP2 s*equences encoding the transit peptide and the 3'-UTR were obtained from GenBank accession NCAA01008649.

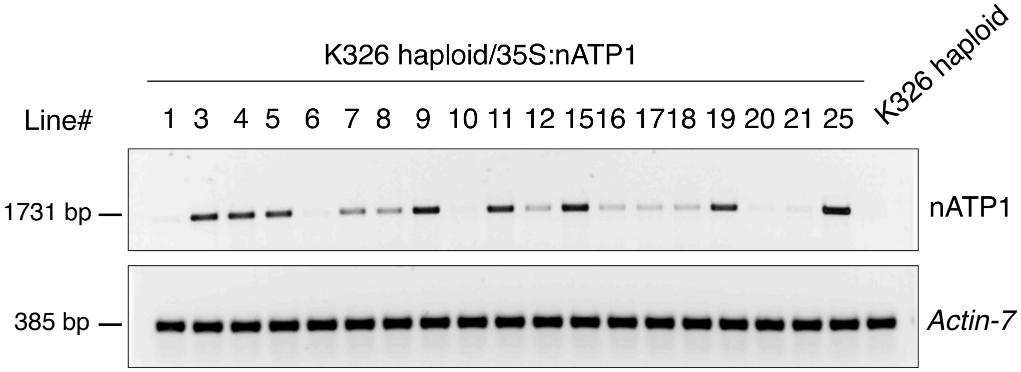


**Figure S2. Semi-quantitative PCR analysis of K326 haploid plants transformed with 35S:nATP1.** The tobacco *Actin-7* gene (GenBank accession XM_016658880) was used as a control. PCR reactions were conducted for 18 cycles and 21 cycles for nATP1 and *Actin-7*, respectively.

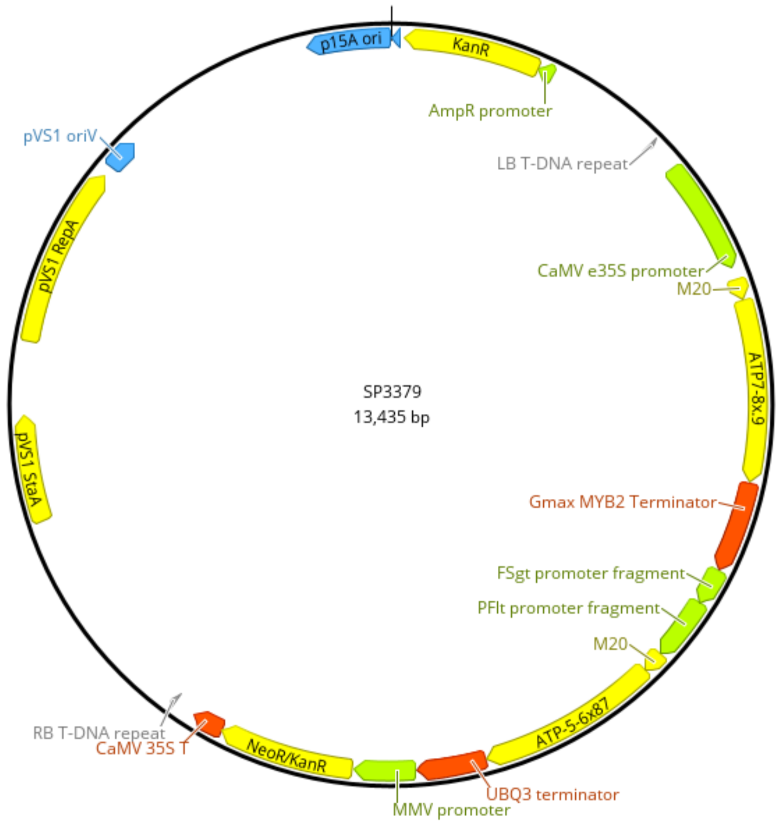


**Figure S3. MitoARCUS vector SP3379.** ATP 5-6x87 and ATP 7-8x.9 encode ARCUS enzymes designed to target two distinct 22 bp sequences in *atp1*. M20, transit peptide from putative Arabidopsis HNH endonuclease (GenBank accession KAG7646785). ATP 5-6x87 is under the transcriptional control of a chimeric *Figwort Mosaic Virus – Peanut Chlorotic Streak Caulimovirus* promoter; termination of transcription is mediated by the termination sequences of the soybean *Ubiquitin3* (UBQ3) gene. ATP 7-8x.9 is under the transcriptional control of an enhanced CaMV 35S promoter; termination of transcription is mediated by the termination sequences of the soybean *MYB2* gene. Expression of the *nptII* selectable marker gene (NeoR/KanR) is driven by a *Mirabilis Mosaic Virus* (MMV) promoter and terminated by the CaMV 35S terminator region. RB, right border; LB, left border.

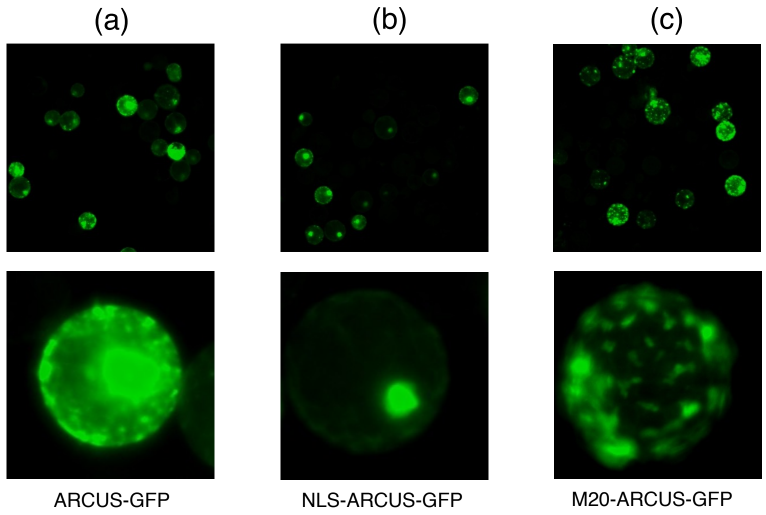


**Figure S4. Transient expression of ARCUS-GFP fusion proteins in tobacco protoplasts.** ARCUS-GFP fusions were introduced with: (a) no localization peptide; (b) a nuclear localization signal (NLS); and (c) the transit peptide M20 from a putative Arabidopsis HNH endonuclease (GenBank accession KAG7646785). Pictures were taken at 20X and 40X magnification (top and bottom, respectively). M20 transit peptide sequence: MGSSFSASFTNSTTAAAVPPPSPPSSPSRSNVKSNGEERPRF.

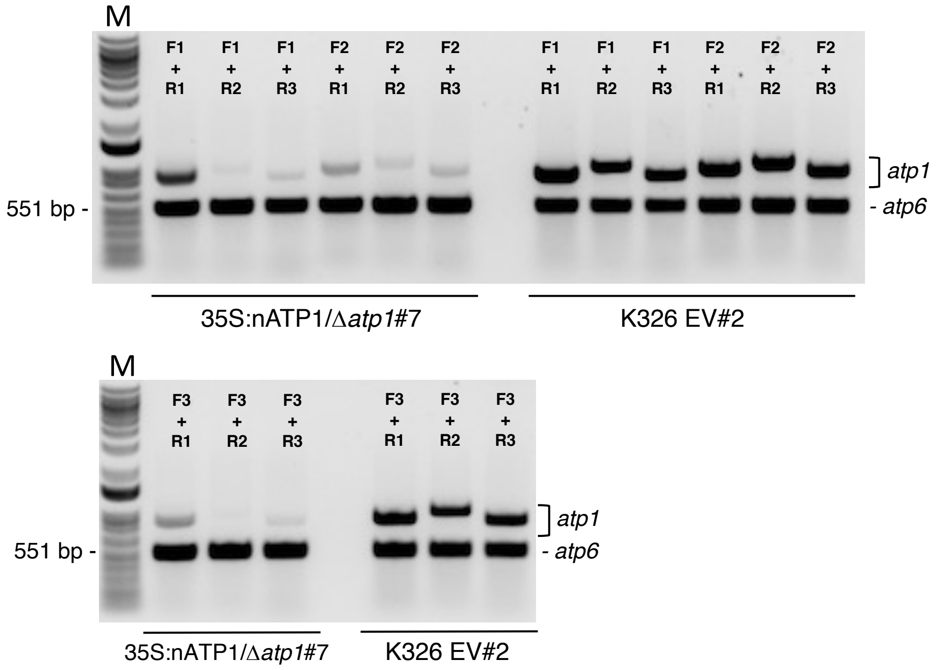


**Figure S5. Primer combinations designed to amplify *atp1.*** Primer sequences and sizes of predicted amplification products are shown in Table S1. F1 = atp1_F1; F2 = atp1_F2; F3 = atp1_F3; R1 = atp1_R1; R2 = atp1_R2; R3 = atp1_R3. M, molecular weight marker. EV, empty vector.

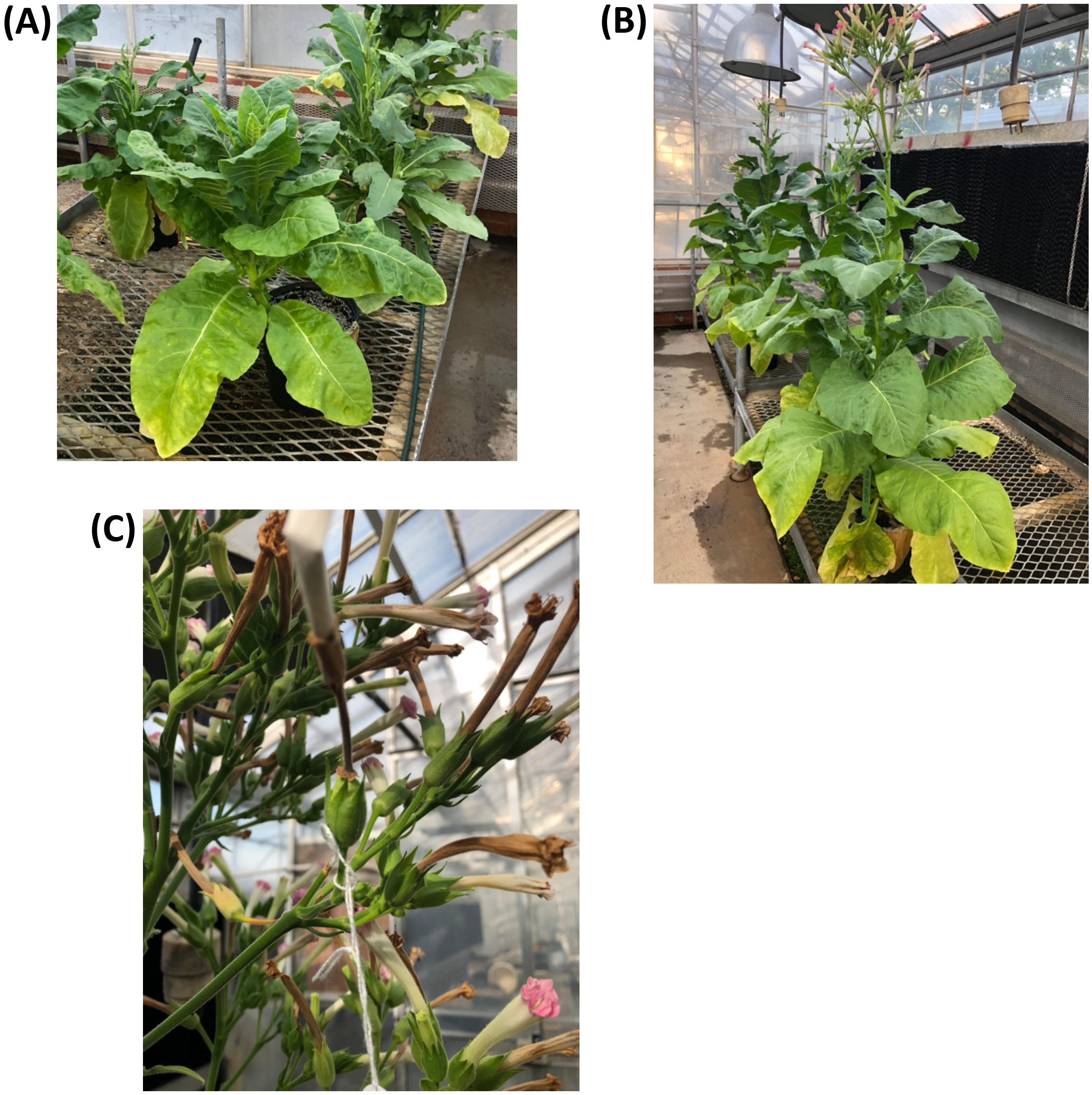


**Figure S6. Examples of mature $T_0$ 35S:nATP1/Δ*atp1* plants.** Representative 35S:nATP1/Δ*atp1* plants that display abnormal **(A)** or typical **(B)** growth patterns are shown 10 weeks after transplanting to soil. **(C)** Ovary/capsule development is observed in flowers that have been cross-pollinated. Fertilized flower has a paper pollination tube and is tagged with a string at the base.

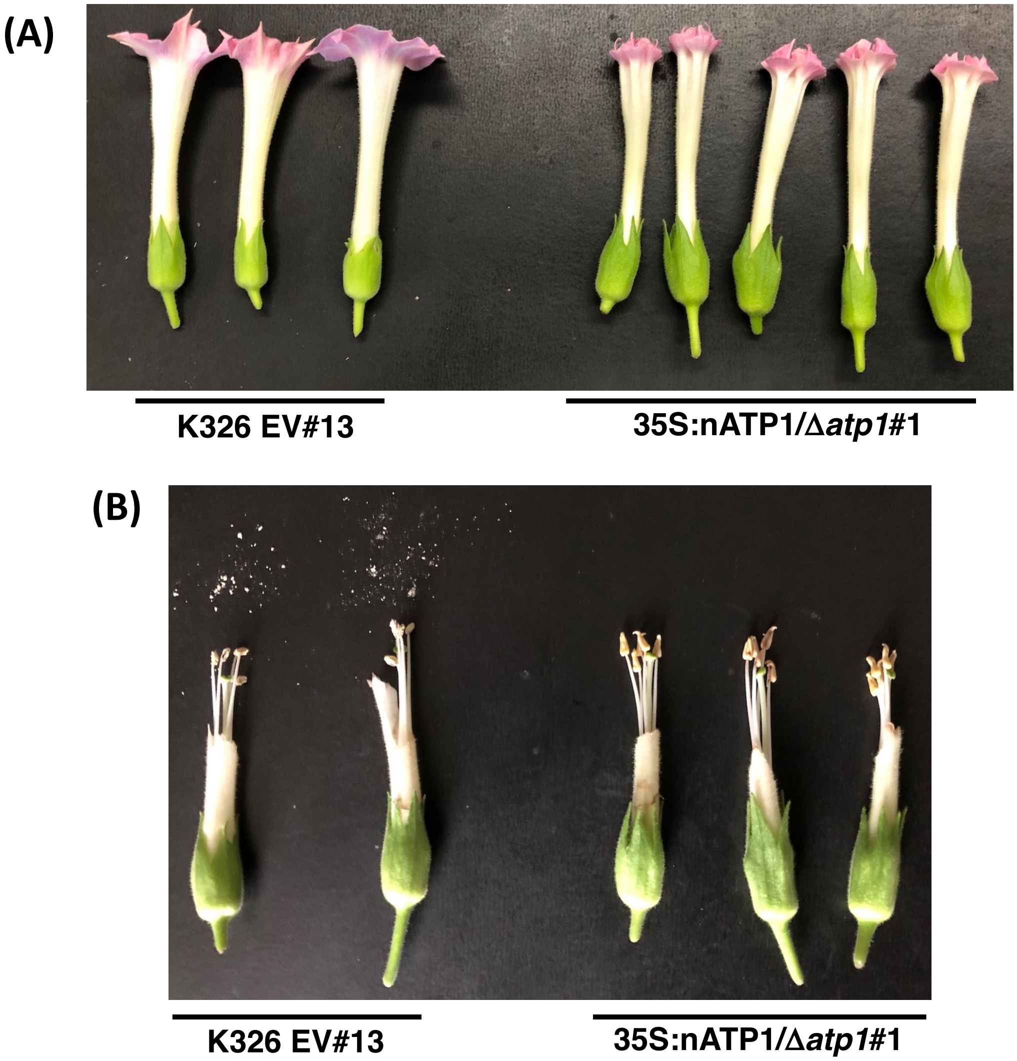


**Figure S7. Examples of normal and 35S:nATP1/Δ*atp1* flowers and anthers.** Empty vector control (K326 EV#13) and 35S:nATP1/Δ*atp1*#1 whole flowers **(A)** and anthers **(B)** are shown at Stage 12, when dehiscence normally occurs. Prior to being photographed, tobacco anthers were tapped on the black background to reveal the presence or absence of pollen.

**(A)**

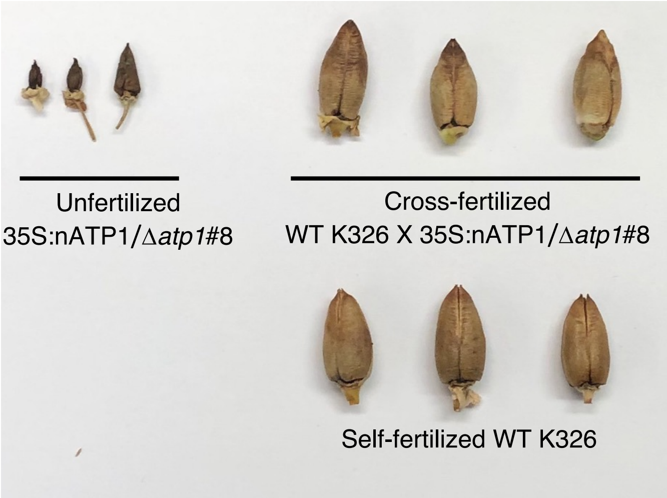


**(B)**

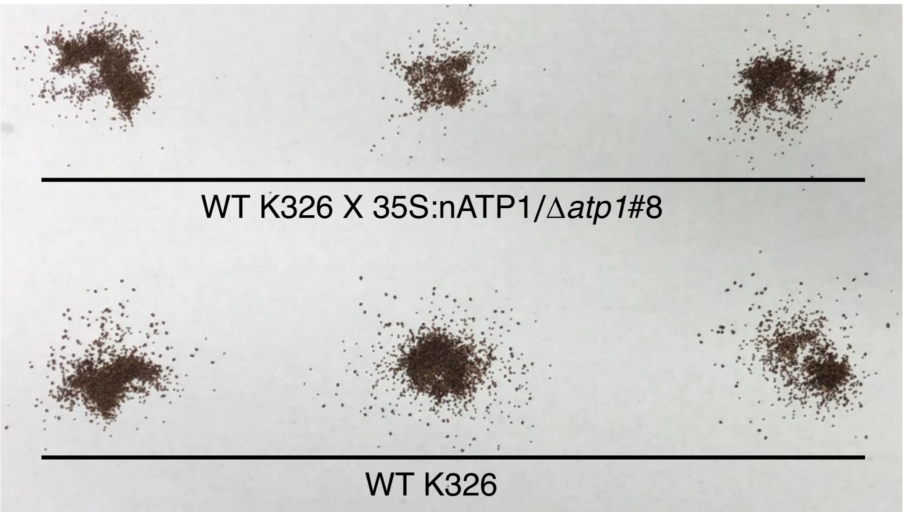


**Figure S8. Mature capsules (A) and seeds (B) of plant 35S:nATP1/Δ*atp1*#8 fertilized with WT tobacco pollen.**

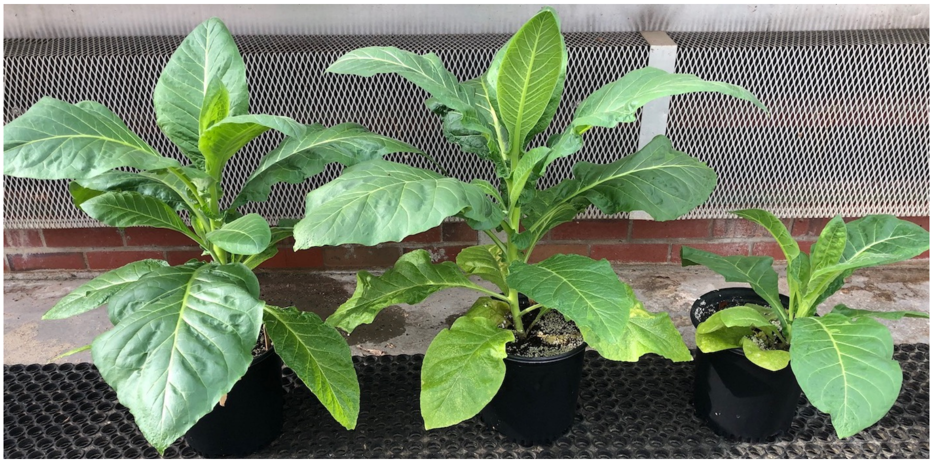


**Figure S9. Plants used for PacBio sequence analysis.** Tissue culture [tc] was used to propagate $T_0$ events 35S:nATP1/Δ*atp1*#22, #8 and #16. Plants derived from the original $T_0$ event 35S:nATP1/Δ*atp1*#16 through culturing displayed a stunted growth phenotype similar to the parental plant.

**(A)**

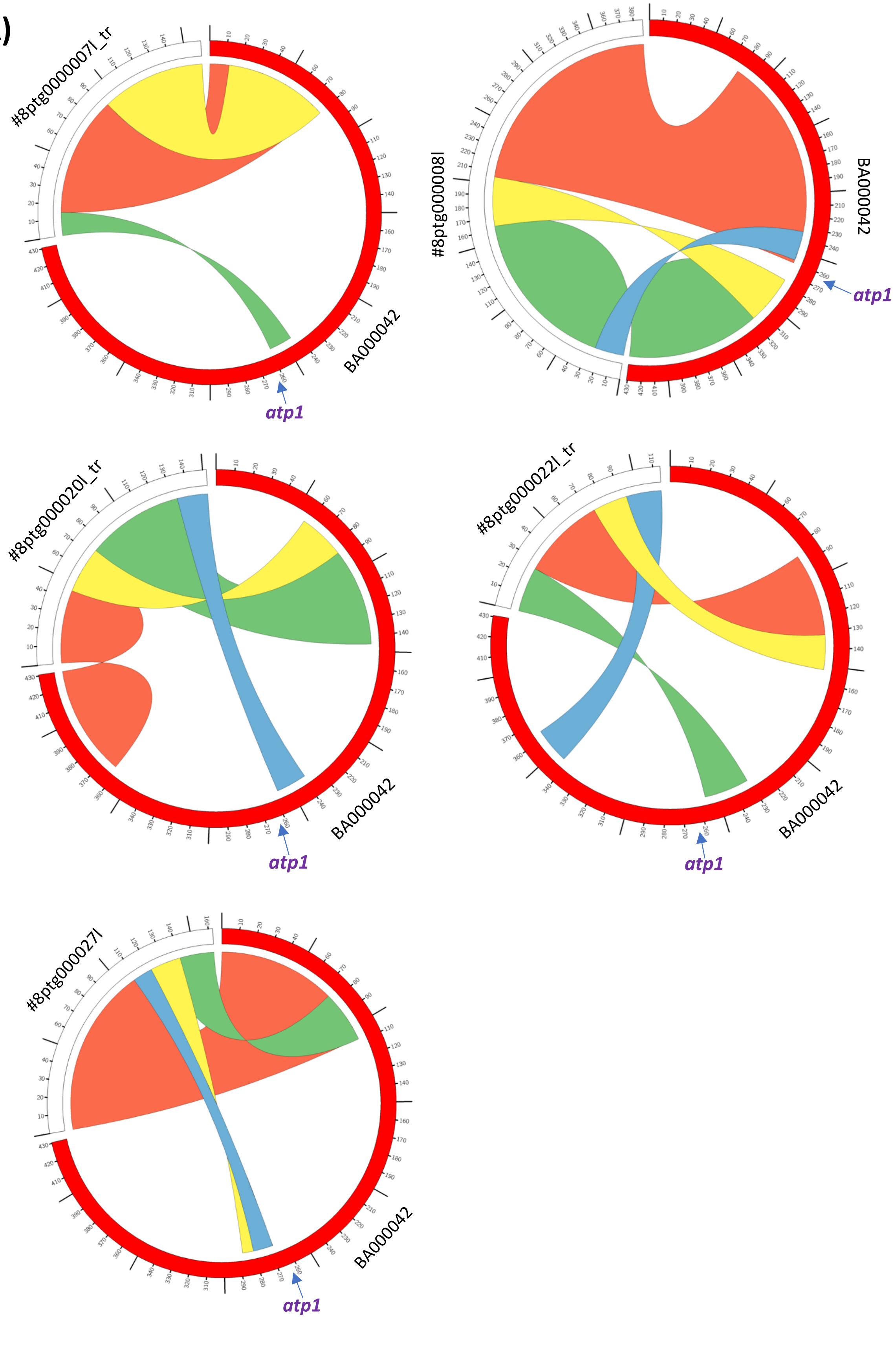

#8ptg000007l_tr
BA000042
atp1
#8ptg000008l
BA000042
atp1
#8ptg000020l_tr
BA000042
atp1
#8ptg000022l_tr
BA000042
atp1
#8ptg000027l
BA000042
atp1

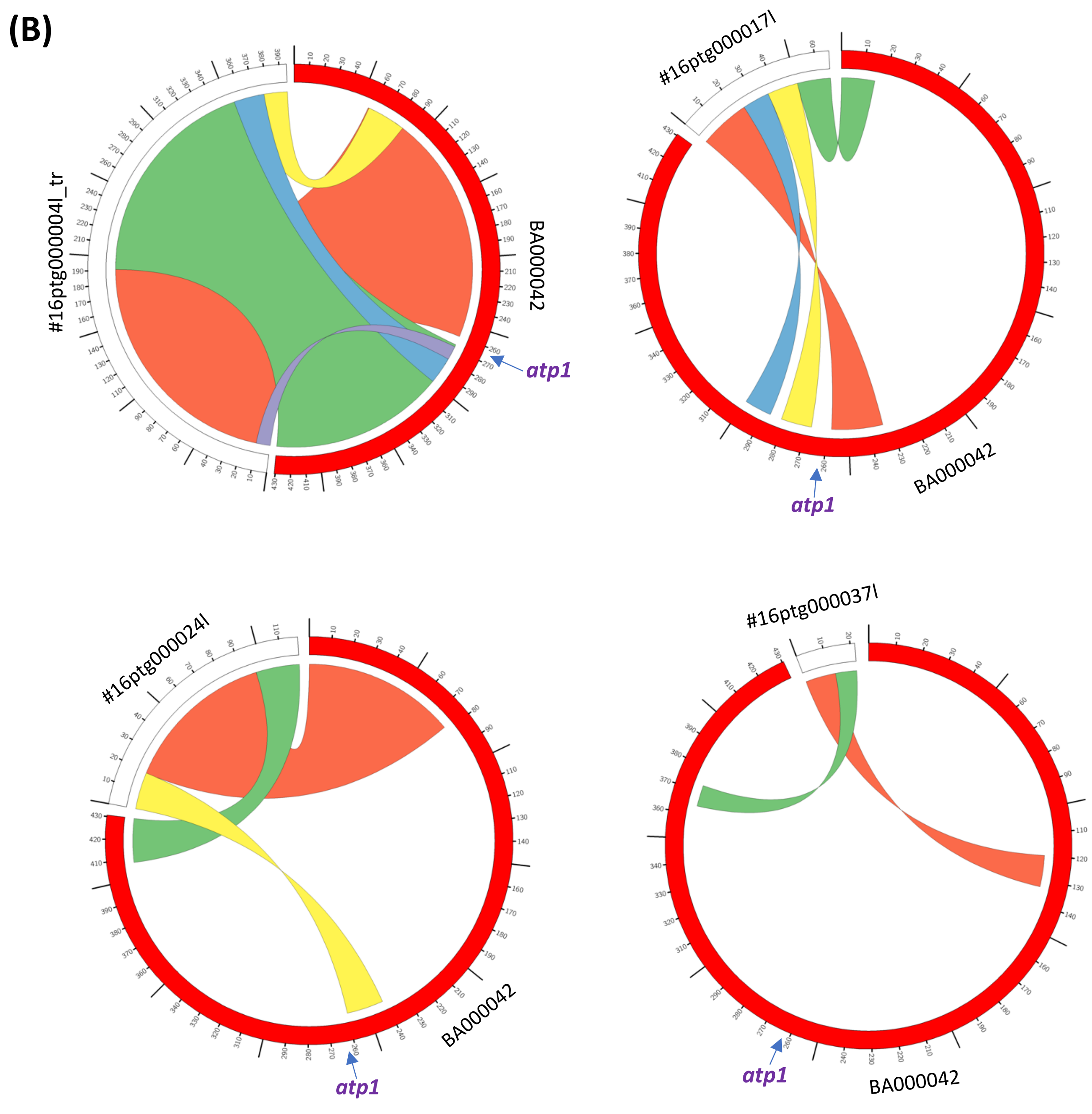
(B)
#16ptg000004l_tr
BA000042
atp1
#16ptg000017l
BA000042
atp1
#16ptg000024l
BA000042
atp1
#16ptg000037l
BA000042
atp1

(C)

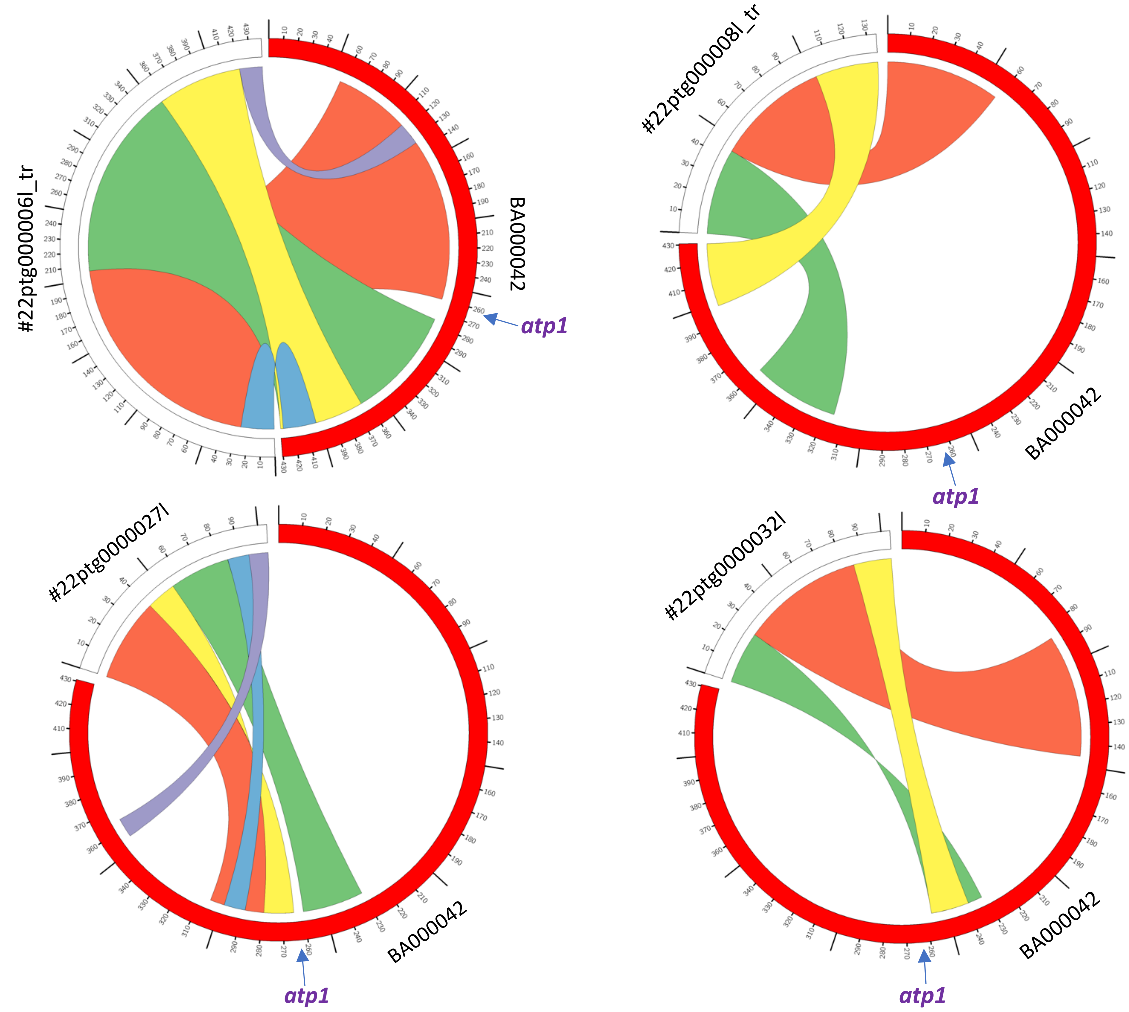


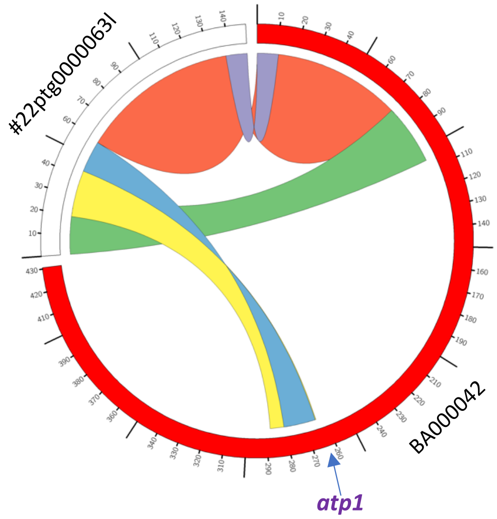


**Figure S10. Circos plots of contigs corresponding to the mitochondrial genomes of 35S:nATP1/Δ*atp1*#8 (A), 35S:nATP1/Δ*atp1*#16 (B), and 35S:nATP1/Δ*atp1*#22 (C) aligned to the tobacco mitochondrial reference genome BA000042.** Blocks of sequence that align between the contigs and reference genome are connected by arcs of the same color. Contigs that end with ‘tr’ were manually trimmed to remove NUMT sequences located at the 5’ or 3’ ends. The location of *atp1* on the reference genome is indicated on each plot.

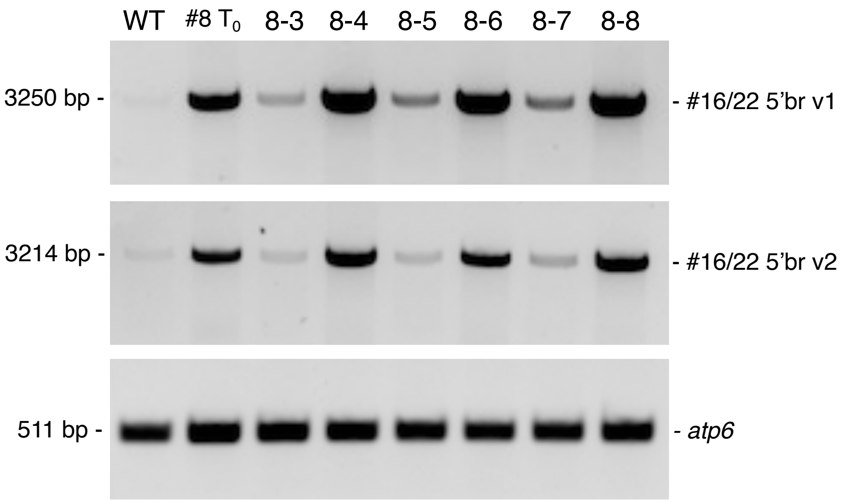


**Figure S11. SSS in 35S:nATP1 X 35S:nATP1/Δ*atp1*#8 $F_1$ progeny.** Semi-quantitative PCR was conducted using primers specific for the two subgenomic variants shown in Figure 6c. The tobacco *atp6* gene was used as a control. All PCR reactions were run for 25 cycles.

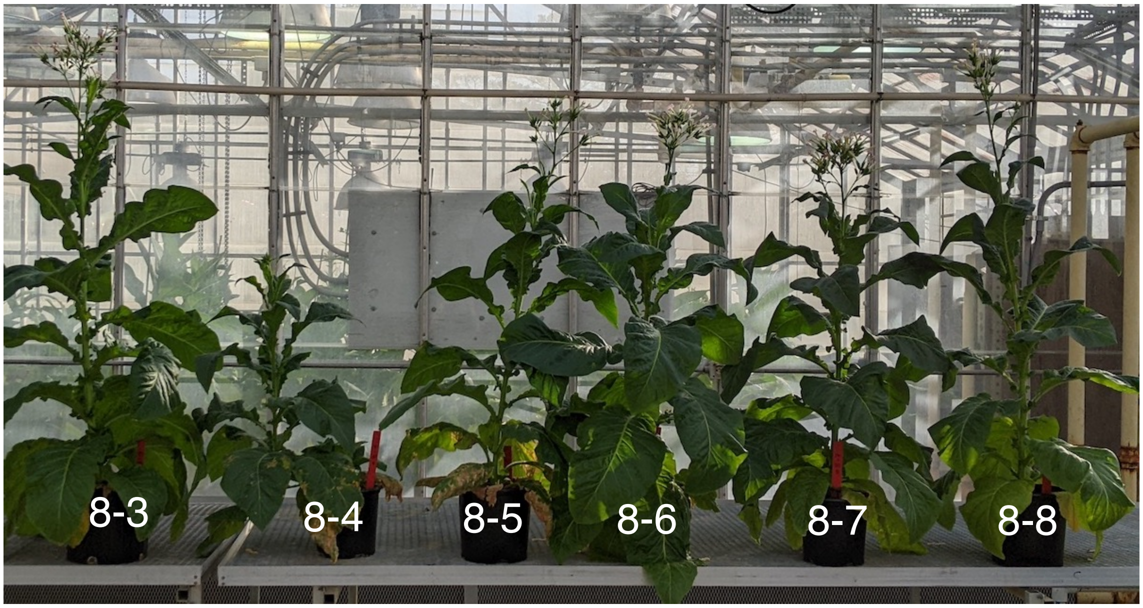


**Figure S12. 35S:nATP1 X 35S:nATP1/Δ*atp1*#8 $F_1$ plants at flowering.**
