## Supplementary material for "Cytoplasmic Male Sterility and Abortive Seed Traits Generated through Mitochondrial Genome Editing Coupled with Allotopic Expression of *atp1* in Tobacco": Table S1

### Table S1. PCR primers and reaction conditions used in this study

| Name | Sequence (5'-3') | Purpose | Product Size | Annealing Temp/time | Extension Temp/time |
|---|---|---|---|---|---|
| ATP2TP_F<br>ATP1_R | ACTATGGCTTCTCGGAGGCTTCTC<br>TCAGATGAATGCGAGTGCGGA | Semi-quantitative PCR of 35S:nATP1 $T_0$ individuals | 1731bp | 65°C/30 sec | 72°C/2 min |
| NtAct_F4<br>NtAct_R2 | AGCACCCTGTTCTTCTCAC<br>GTCAAGCTCCTGCTCGTAG | *Actin7* amplification | 385bp | 59°C/30 sec | 72°C/40 sec |
| atp1_F1<br>atp1_R1 | TGAGATCGGTCGAGTGGTCT<br>TACGTCTCCAGCCTGTGTTT | *atp1* amplification | 928bp | 64°C/15 sec | 72°C/30 sec |
| atp1_F1<br>atp1_R2 | TGAGATCGGTCGAGTGGTCT<br>ACTGACAGATAAGCCGACGT | *atp1* amplification | 1042bp | 64°C/15 sec | 72°C/30 sec |
| atp1_F1<br>atp1_R3 | TGAGATCGGTCGAGTGGTCT<br>TAAGGCGGTCAAGCTACCTG | *atp1* amplification | 898bp | 63°C/15 sec | 72°C/30 sec |
| atp1_F2<br>atp1_R1 | GGAACTTTCTCCCCGAGCTG<br>TACGTCTCCAGCCTGTGTTT | *atp1* amplification | 1011bp | 64°C/15 sec | 72°C/30 sec |
| atp1_F2<br>atp1_R2 | GGAACTTTCTCCCCGAGCTG<br>ACTGACAGATAAGCCGACGT | *atp1* amplification | 1125bp | 64°C/15 sec | 72°C/30 sec |
| atp1_F2<br>atp1_R3 | GGAACTTTCTCCCCGAGCTG<br>TAAGGCGGTCAAGCTACCTG | *atp1* amplification | 980bp | 65°C/15 sec | 72°C/30 sec |
| atp1_F3<br>atp1_R1 | CAAGTGGATGAGATCGGTCG<br>TACGTCTCCAGCCTGTGTTT | *atp1* amplification | 936bp | 63°C/15 sec | 72°C/30 sec |
| atp1_F3<br>atp1_R2 | CAAGTGGATGAGATCGGTCG<br>ACTGACAGATAAGCCGACGT | *atp1* amplification | 1050 bp | 63°C/15 sec | 72°C/30 sec |
| atp1_F3<br>atp1_R3 | CAAGTGGATGAGATCGGTCG<br>TAAGGCGGTCAAGCTACCTG | *atp1* amplification | 906bp | 63°C/15 sec | 72°C/30 sec |
| atp6_F1<br>atp6_R3 | ATCCGGGCCTTAATCCTTGC<br>TGCGAGGGGAAAACTTTTGT | *atp6* amplification | 511 bp | varied | varied |
| hptII_F<br>hptII_R | GTGTACGCCCGACAGTCCCGGC<br>CCCGATTCCGGAAGTGCTTGAC | *HPTII* amplification | 705bp | 63°C/20 sec | 72°C/15 sec |
| nptII_F<br>nptII_R | GAACAAGATGGATTGCACGC<br>AGAAGGCGATAGAAGGCGAT | *NPTII* amplification | 773 bp | 63°C/15 sec | 72°C/15 sec |
| E10_F<br>E10_R | CAGCAGATAGTGACGCTAACC<br>AACCCTGGATCAAGTGTGCG | *CYP82E10* amplification | 549 bp | 63°C/15 sec | 72°C/15 sec |
| ARCUS_F1<br>ARCUS_R1 | CCAACCACGTCTTCAAAGCA<br>ATAGACCTCCCACGGATCCT | Amplification of SP3379 construct | 714 bp | 64°C/15 sec | 72°C/30 sec |
| #8 5'br_F3<br>#8 5'br_R4 | ATCGAACTTGAACTCCGGGT<br>CCAACAACTTCACTCCGGAG | Amplification of recombination junction at 5' break point in event 35S:nATP1/Δ*atp1*#8 | 843 bp | 63°C/15 sec | 72°C/30 sec |
| #8 3'br_F2<br>#8 3'br_R3 | CCCTCAAAGCCCCATTAACG<br>AAGACCTAAGCTTGAAGTCAAG | Amplification of recombination junction at 3' break point in event 35S:nATP1/Δ*atp1*#8 | 754 bp | 61°C/15 sec | 72°C/30 sec |
| #16/22 5'br:v1_F3<br>#16/22 5'br:v1_R1 | ATCGAACTTGAACTCCGGGT<br>ATCTTCACTTTACGCCACGC | Amplification of recombination junction at 5' break point in event 35S:nATP1/Δ*atp1*#8 and #22 (variation 1) | 3250 bp | 64°C/20 sec | 72°C/2 min |

| | | | | | |
|---|---|---|---|---|---|
| #16/22 5'br:v2_F5<br>#16/22 5'br:v2_R4 | AGCGCAGTACTTCCGTAACT<br>TATCGCCCTTTGTTCCTCCA | Amplification of recombination junction at 5' break point in event 35S:nATP1/Δ*atp1*#8 and #22 (variation 2) | 3214 bp | 64°C/20 sec | 72°C/2 min |
| #16 3'br_F1<br>#16 3'br_R2 | TATGGATAGGGCGACGTGAC<br>GCCAAGACTGTACGAGGAGA | Amplification of recombination junction at 3' break point in event 35S:nATP1/Δ*atp1*#16 | 734bp | 64°C/15 sec | 72°C/30 sec |
| #22 3'br_F1<br>#22 3'br_R2 | GTTCGGTCAGGCTTGCTTAG<br>AGTAAGGCATTGGGGATCGT | Amplification of recombination junction at 3' break point in event 35S:nATP1/Δ*atp1*#22 | 790 bp | 64°C/15 sec | 72°C/30 sec |

- All reactions were initiated with a 30 sec denaturation at 98°C for 30 sec, and terminated with a 7 min extension at 72°C after 33 cycles (or number of indicated cycles for semi-quantitative PCR assays).
- Reactions were conducted using Phusion Taq and 5x HF reaction buffer (New England Biolabs – cat. #M0530L).
- All denaturation steps were conducted at 98°C for 10 sec.
- Unless otherwise indicated, primers were added at a concentration of 10 μM.
- The *atp6* primers were amplified using the conditions preferred for amplification of the *atp1* primers they were paired with.

### Table S2. PacBio sequence assembly summary

| Line name | Total reads | N50 of Reads | Reads that map to Mt genome | % Reads mapping to Mt genome | Number of contigs in assembly* | Number of contigs that represent NUMTs* | Number of legitimate Mt genome contigs** |
|---|---|---|---|---|---|---|---|
| 35S:nATP1 Δ*atp1*#8 | 878,706 | 14,113 | 40,862 | 4.7% | 94 | 89 | 5 |
| 35S:nATP1 Δ*atp1*#16 | 543,041 | 14,286 | 18,282 | 3.4% | 40 | 36 | 4 |
| 35S:nATP1 Δ*atp1*#22 | 911,783 | 13,168 | 44,784 | 4.9% | 95 | 90 | 5 |

*In addition to the mitochondrial genome, this includes contigs that share homology to the chloroplast genome, as the mitochondrial genome also contains fragments of the chloroplast genome

**A contig was classified as legitimate if the sequence displayed >99.99% nucleotide identity to the tobacco mitochondrial reference genome (BA000042) over its entire length, and an NUMT if it shared <98% identity to the reference genome over its length.

Mt, mitochondrial
